# Dietary lipid is largely deposited in skin and rapidly affects insulating properties

**DOI:** 10.1101/2024.10.03.615679

**Authors:** Nick Riley, Ildiko Kasza, Isabel D.K. Hermsmeyer, Michaela E. Trautman, Greg Barrett-Wilt, Raghav Jain, Judith A. Simcox, Chi-Liang E. Yen, Ormond A. MacDougald, Dudley W. Lamming, Caroline M. Alexander

**Affiliations:** McArdle Laboratory for Cancer Research, University of Wisconsin-Madison; Department of Molecular & Integrative Physiology, University of Michigan; Department of Medicine, University of Wisconsin-Madison; William S. Middleton Memorial Veterans Hospital, Madison; Biochemistry Mass Spectrometry Core; Department of Biochemistry, University of Wisconsin-Madison; Howard Hughes Medical Institute, University of Wisconsin-Madison; Department of Nutritional Sciences, University of Wisconsin-Madison

## Abstract

Skin has been shown to be a regulatory hub for energy expenditure and metabolism: mutations of skin lipid metabolism enzymes can change the rate of thermogenesis and susceptibility to diet-induced obesity. However, little is known about the physiological basis for this function. Here we show that the thermal properties of skin are highly reactive to diet: within three days, a high fat diet reduces heat transfer through skin. In contrast, a dietary manipulation that prevents obesity accelerates energy loss through skins. We found that skin was the largest target in a mouse body for dietary fat delivery, and that dietary triglyceride was assimilated both by epidermis and by dermal white adipose tissue. Skin from mice calorie-restricted for 3 weeks did not take up circulating lipids and showed a highly depleted stratum corneum. Dietary triglyceride acyl groups persist in skin for weeks after feeding. Using multi-modal lipid profiling, we have implicated both keratinocytes and sebocytes in the altered lipids which correlate with thermal function. In response to high fat feeding, wax diesters and ceramides accumulate, and triglycerides become more saturated. In contrast, in response to the dramatic loss of adipose tissue that accompanies restriction of the branched chain amino acid isoleucine, skin becomes more heat-permeable, resisting changes induced by Western diet feeding, with a signature of depleted signaling lipids. We propose that skin should be routinely included in physiological studies of lipid metabolism, given the size of the skin lipid reservoir and its adaptable functionality.

## INTRODUCTION

A growing body of research highlights the influence of altered skin function on system-wide metabolism and energy expenditure. Mice with gene mutations that specifically affect the lipid composition of their skin exhibit changes in thermoregulation, resistance to diet-induced obesity, and enhanced insulin sensitivity^1,2^. As one example, mice with a keratinocyte-specific mutation of the lipid transport protein, acyl-CoA-binding protein (ACBP), show impaired thermal barrier function (increased TEWL), increased energy expenditure and food intake, altered lipid metabolism in liver, increased lipolytic flux in white adipose tissue (WAT), iWAT beiging, and resistance to diet-induced obesity. These systemic changes of energy homeostasis are reversed by housing mice at thermoneutrality or inhibiting thermogenic β-adrenergic signaling^3,4^. Other examples of epidermal regulation of energy homeostasis include genetic studies of ACER1, DGATs, ELOVL and SCD enzymes, discussed in detail by us and others^2,5–9^. These studies conclude that the rate of heat loss through skin determines the rate of heat production required to maintain body temperature, which in turn regulates energy homeostasis, to determine the overall metabolic strategy.

However, the mechanisms underlying the regulation of the skin thermal barrier are not known, and this presents a significant translational knowledge gap, as the manipulation of skin properties may offer a valuable approach for improving health. Skin is comprised of at least three, closely-apposed lipid-enriched layers: these are the pelt (hair), with associated waxy sebome, the stratum corneum, assembled by epidermal keratinocytes, and a skin-associated adipose depot, the dermal white adipose tissue (dWAT), with unique function and regulation^5,10^. The properties of any of the three biomaterials could therefore be important effectors of energy expenditure.

In this study, we test whether total skin function is an early responder to conditions that change metabolism, and if so, which of the cell types and thermal components correlate with functional changes. We focus on the molecular lipidome, for two reasons. The first is the body of experimental data demonstrating that mutations of epidermal lipid metabolic enzymes confer changes of system-wide metabolism, and the second is that lipid composition is known to be modified by environmental temperature changes throughout the biological phyla, suggesting that altered lipid biosynthesis is an ancient means of temperature-induced adaptation^11^.

Using a short timeline of 3 days, we find that skins become more insulating in response to an obesogenic high fat diet. Indeed, we find the major site of delivery for dietary fats is mouse skin, and delivery is ablated if mice are calorie-restricted. Vice versa, we find that skins become more heat-permeable in response to a diet low in the branched chain amino acid, isoleucine, which is a diet that promotes lean-ness, health and longevity. Overall, we conclude that skin properties are highly modifiable, and amongst the earliest functional responders to diets that affect obesogenesis and energy expenditure. Using multi-modal lipidomics, we show that the amount and specifics of the lipids from sebome and epidermis change in parallel with functional changes, suggesting that both sebocyte and keratinocyte could be manipulated to promote heat flux through skins.

## RESULTS

### Skin is an early responder to a change of diet

Most studies of metabolic adaptation to diet or environment use a minimum 2-3 weeks after the diet switch before testing for phenotypic alterations of energy expenditure. Our goal was to evaluate whether skin adaptation preceded or followed other significant changes, such as an increased body weight in response to high fat feeding. We therefore chose 3 days after the dietary switch to a high fat diet (**HFD**; 60% calories from fat, provided as 90% lard, 10% soybean oil, compared to control diet 18% calories from fat, provided as soybean oil) for subsequent analyses.

Both male and female BALB/cJ mice ate more kilocalories (Kcal) per day when switched to HFD (**Fig.1A**); however, these mice did not yet show a significant change of body weight within 3 days (**Fig.1B**). The average area of perigonadal white adipocytes and dermal white adipocytes was also unchanged and there were no significant changes of skin morphology, including dWAT thickness (**Fig.S1**). Male and female skins are highly dimorphic; for example, male skins (mice under 6 months, fed chow) have much less dWAT than females, with thick collagenous dermis (examples of each are shown in **Fig.1C**).

**Fig. 1.**
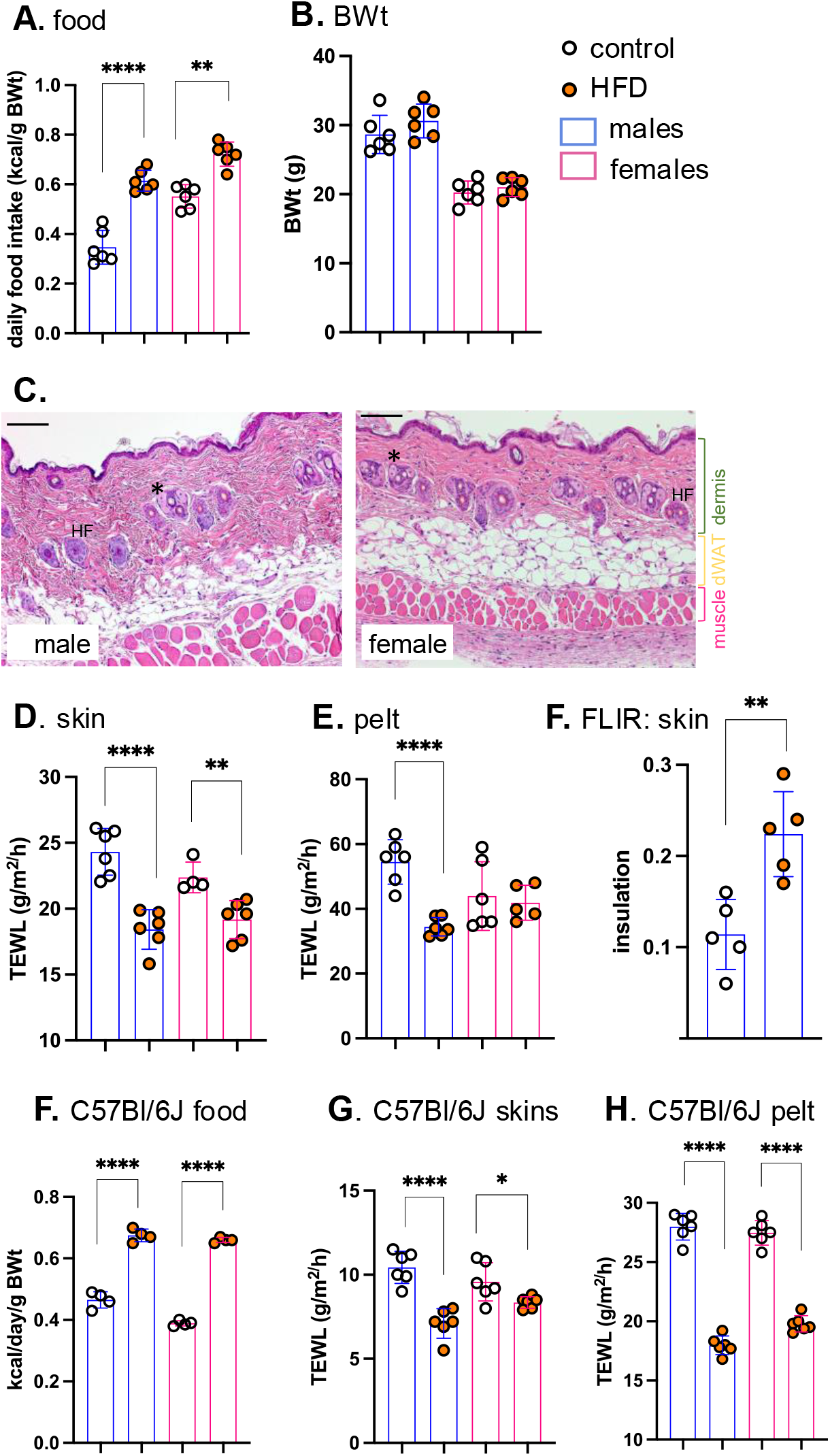
High fat diet consumption rapidly results in lower heat loss through skins. Male and female BALBc/J mice (10-15 weeks old) were switched to HFD for 3 days, and daily food consumption assessed together with their body weight (BWt; n=6; **A,B**). **C**. Representative H&E-stained skin sections are shown for male and female BALB/cJ mice (see also Fig.S1 for comparison of skins from mice fed HFD for 3 days). **\*** marks sebaceous glands, filled with sebocytes. Distinct tissue layers are marked. Scale bar=100μm. Skin thermal properties were measured by TEWL assay of skin (**D**) and pelt (**E;** n=6), and by FLIR assay of relative surface temperature, where ^0^C below adjacent heat block is used to indicate relative insulation (**F;** n=5). To establish this response for C57BL/6J mice, males and females were fed HFD for 3 days and assayed for their relative food consumption (**F;** n=3-4), and the properties of their skins (assayed by TEWL of skin and pelt; **G,H;** n=6). * p<0.05; ** p<0.01, *** p<0.001, **** p<0.0001.

To assess the effect of HFD on the thermal properties of skins, two techniques were employed. For the first, we rely on the assay of trans-epidermal water loss (**TEWL**), a marker of rates of evaporative cooling. We showed previously that evaporative cooling is likely to be a major effector of energy loss for mice^6^; this is “insensible” water loss at environmental temperatures below body temperature, not associated with the heat loss mechanisms activated when body temperature is too high, or the mouse is stressed (such as superficial blood vessel dilation). This assay is done *ex vivo* on a warm, wet heat block, to eliminate effects attributable to changing blood flow, and reported for shaved skin (dorsal “**skin**”) and **pelt** (unshaved), as described previously^6^. Heat transfer across HFD-fed mouse skin and pelt was significantly reduced, notably for males (**Fig.1D, E**). Surface thermography using a forward-looking, infrared camera (**FLIR**) was used to assess radiative heat losses and skin insulation properties. HFD-fed mice showed a 2-fold increase in insulation (**Fig.1F**). We also assessed C57BL/6J mice as the most common model of obesogenesis^12^, and found that in both male and female mice, there was much reduced heat loss across skins and pelts within 3 days of high-fat feeding (**Fig.1F-H**).

### A large fraction of dietary lipid is taken up by and stored in skin

At least two possibilities could explain a rapid response of skin to diet: the first is that high fat consumption sends an indirect cue to skin, and the second is that incorporation of dietary fats into skins changes thermal properties. To address these possibilities, we first tested whether lipids could be delivered directly to skin from high fat diet. We used the approach to studying uptake of dietary lipids that was described by Bartelt et al^13^. We administered ^3^H-[9,10] triolein by gavage (**Fig.2**,**3**), and assessed the assimilation of radiotracer over the course of 2 weeks in serum, liver, BAT, iWAT, pgWAT and skins, for BALB/cJ males and females housed at either thermoneutrality (29^0^C; TMN) or at 10^0^C (cool).

**Fig. 2.**
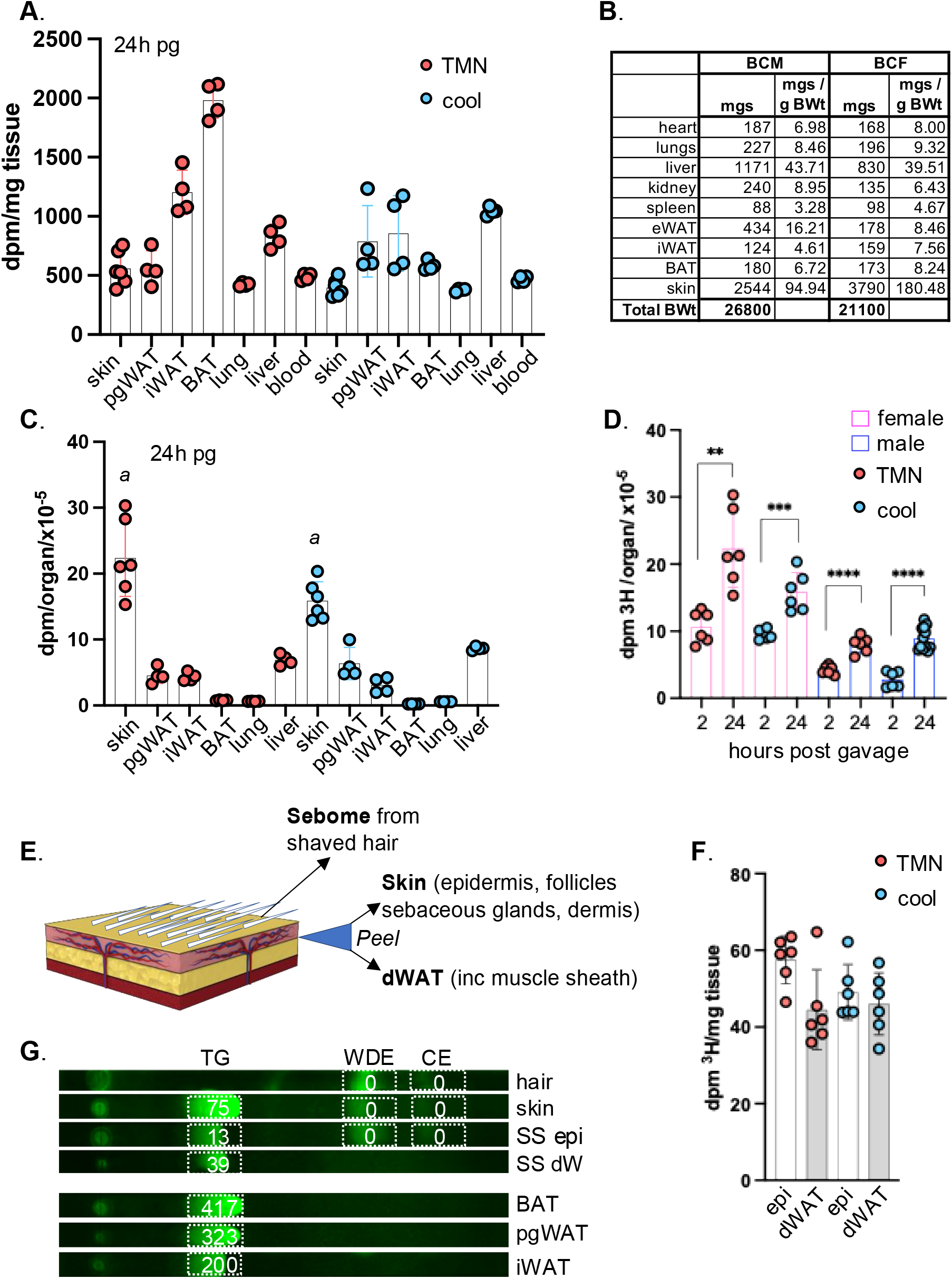
Skins assimilate more dietary acyl lipid than any other organ. BALBc/J female mice were administered 3H-triolein by gavage and assessed 24 hours later (**24h** post-gavage, **pg**) for incorporation of radiolabel into the tissues indicated (pgWAT, perigonadal (visceral) WAT; iWAT, inguinal (subcutaneous) WAT; BAT, brown adipose tissue). Mice were housed either at thermoneutrality (TMN, 29^0^C) or cool (10^0^C); n≥3. **A**. Radiolabel assimilation was calculated as dpm/mg tissue for the tissues indicated. **B.** Average size of each organ was measured and expressed as mgs/g BWt. **C.** Calculated per organ, skins assimilated more label than any other tissue. Assimilation into iWAT was measured for the pair of iWAT depots, noting that there are 10 fat pads altogether in this class. With thermogenic fatty acid oxidation active (cool conditions), skin specific activity was reduced by 30%: ***a***, comparison of skin from mice housed at TMN versus cool, p=0.0001. **D**. Skin from BALB/c female and males 2 and 24 hours after radiolabel administration showed a progressive increase in specific activity; n≥3. **E-G**. Skins from BALB/cJ females were separated into 3 fractions, **sebome** (extracted from hair), **dWAT** (separated from skin), and the skin remainder (**epi**). **F.** Specific activities of epi and dWAT fractions from BALBc/J females housed at TMN or 10^0^C (n=3) showed approximately equal distribution. **G.** Lipid classes were separated by thin layer chromatography (TLC), into triglyceride (TG), wax diester (WDE) and cholesterol ester (CE) and scraped for counting (a representative image is labeled with dpm/mg of tissue equivalent), from 3 adipose depots from BALB/cJ females, along with hair, skin and separated skins (SS). All radiolabel was retrieved from the triglyceride fraction.

### Along with other adipose depots, the lipid oxidation associated with thermogenesis depletes skin

*TGs*. Like other adipose depots, skin showed rapid and progressive assimilation of dietary ^3^H-triolein during the first 24 hours post-gavage (**Fig.2A-D** and **Fig.3**). Normalized by tissue weight, the specific activity of skin was approximately the same as visceral adipose depots (pgWAT; **Fig.2A**). The well characterized thermogenic depots iWAT and BAT^14^ labeled to a 2-4x higher specific activity than other adipose depots, when mice were housed at thermoneutrality. This illustrates their role as the major metabolic drivers of thermogenesis. As expected, radiolabel was immediately depleted from BAT in mice housed in a cool environment (**Fig.2A-D** and **Fig.3**), where other adipose depots, together with skin, showed substantial radiolabel depletion only 2 weeks later. The specific activity of female skin was 2-3-fold higher than for males (shown in Fig.1C), probably due to relative dWAT content (40% compared to 16% total skin thickness).

**Fig. 3.**
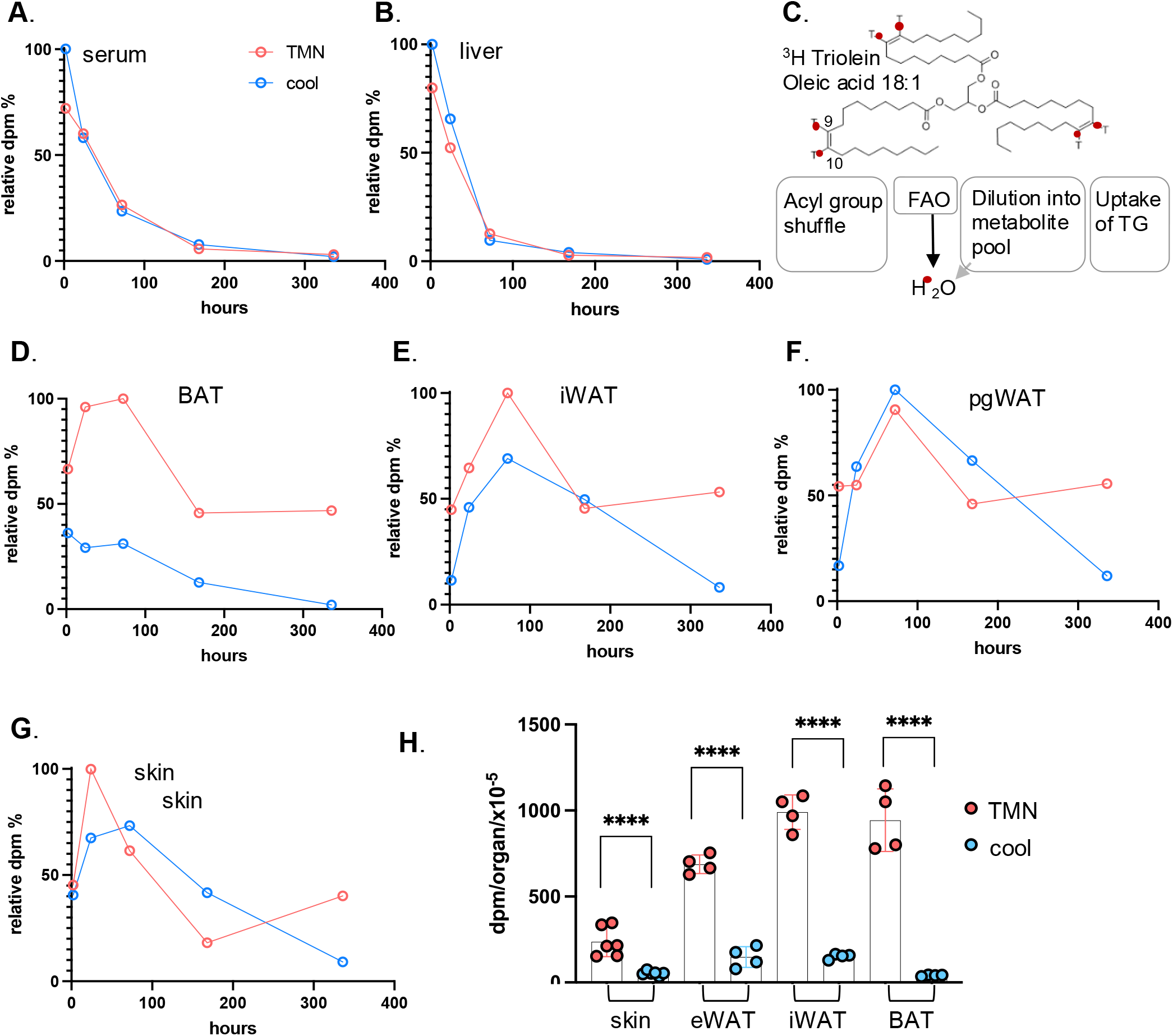
Dietary fat persists in skin and adipose depots for several weeks. **A,B.** Dilution and elimination of the radiolabel in BALB/cJ females is indicated for 2 weeks post-gavage, for serum and liver (see also Fig.S2, dpm/mg tissue, all data points; n≥3). Timepoints shown are 2h, 24h, **3 days** (72h), **1 week** (168h) and **2 weeks** (336h) (n≥3 for each timepoint). **C.** A scheme of the potential fates of ^3^H-triolein (acyl group ^3^H-label): acyl group reshuffling, fatty acid oxidation (FAO), degradation of acyl groups and dilution into general metabolic pool, or uptake of the whole TG moiety. These are not discriminated by assay of dpm/mg tissue. **D-G.** Relative specific activity of adipose depots and skin, expressed with respect to time, for mice housed either at 29^0^C (approximately thermoneutral) or in cool housing (10^0^C). **H.** Statistical analysis of specific activity of tissues indicated for the 2- week timepoint.

When the total incorporation of ^3^H-triolein label is calculated per organ (**Fig.2B**), we found that skin was the largest target for lipid uptake (**Fig.2 B,C**), irrespective of environmental housing temperature. Dedicated lipid storage depots (pgWAT and iWAT) showed only 20% (each) of the total acquisition and storage capacity of skin.

### Dietary TGs are delivered to both epidermis and dWAT

To determine which skin fractions were acquiring dietary lipids, we separated skins into 3 fractions (**Fig.2E**). These reflect the three distinct lipid-based thermal barriers in skin: the pelt/hair, coated in a sebome of mostly wax di-esters; the stratum corneum, a proteo-lipid cross- linked sheath of enucleated keratinocytes providing the top layer of the epidermis; and the dermal white adipose tissue (dWAT) underlying the pelt/epidermis/dermal layers (see **Fig.7**). Using enzymatic dissociation, we separated the dWAT from the dermal/epidermal layer (epi) of shaved skins to generate a highly enriched dWAT adipose fraction (Fig.S10). These fractions (epi and dWAT) were labeled to approximately the same specific activity; 60% recovery from epidermal fraction, and 40% from dermal adipose (**Fig.2F**). We conclude that ^3^H oleate was acquired by the keratinocyte/sebocyte-enriched tissue fraction to a similar extent as the adipose depot. Label accumulation by hair was low/undetectable over the 2-week period.

To determine which lipid class had acquired ^3^H oleate, we isolated the lipids in sebome, skin, and separated skins (SS) by thin layer chromatography (TLC) and found that the triglyceride fraction of skin accounted for all detectable counts (**Fig.2G**). There were no counts in the wax diester (WDE) or cholesterol ester (CE) fraction. Using this method, the TG fraction of BAT yielded the highest specific activity, as expected.

### Dietary TGs persist for weeks in mice housed at thermoneutrality

Predictably, the specific activity of sera and liver diluted out within 4 days (96 hours; **Fig.3 A,B**, **S2**). Measured specific activity (dpm/mg tissue) constitutes an equilibrium between uptake of ^3^H-triolein triglyceride or ^3^H-oleate acyl groups, and the competing oxidation rate of fatty acids, generating ^3^H2O or labeled metabolites (**Fig.3C**). We assume that only intact acyl groups are effectively scored by this labeling strategy.

In contrast to liver and sera, the specific activity of adipose depots and skin did not dilute out rapidly, in fact, when mice were housed at thermoneutrality, the specific activity was maintained to 50% of maximum specific activity even 2 weeks after gavage (**Fig.3D-H**). For mice housed cool, the radiotracer was depleted within 2 weeks, demonstrating that, along with other adipose depots, skin-associated lipids were subject to mobilization and depletion by thermogenic activation. We considered whether the gavage lipid dose could be higher than usual and could be accumulating artifactually in skins. However, the dose administered was only 50% of the lipid intake present in a daily ration of chow.

### Uptake of dietary lipid into skin is suppressed in calorie-restricted mice, with consequences for dWAT and epidermis

We conclude that the rapid and sustained uptake and storage of ^3^H-oleate in skins is consistent with a direct delivery mechanism from diet. We were curious to know whether this delivery could be modified if mice were switched to a “healthy” dietary paradigm. After 3 weeks of classic calorie restriction (**CR**), which is so well known to enhance metabolic health and longevity^15,16^, we found that dWAT of mice was dramatically ablated (>90%). Other depots were also depleted: pgWAT was 80% reduced, iWAT was 60% reduced and overall body weight was reduced by 7.4g (25% total; **Fig.4A-E**). The corresponding uptake of ^3^H-triolein radiotracer into skin-associated TGs was dramatically reduced, by 90% (**Fig.4F**), despite the presence of more label in the circulation (**Fig.4G**). Decreased lipid uptake into skin was associated with dramatically reduced epidermal thickness and mitotic index of the interfollicular basal keratinocytes (**Fig.4H-J**). Together, these data illustrate that lipid delivery to skin is controlled, with profound impact on form and function.

**Fig. 4.**
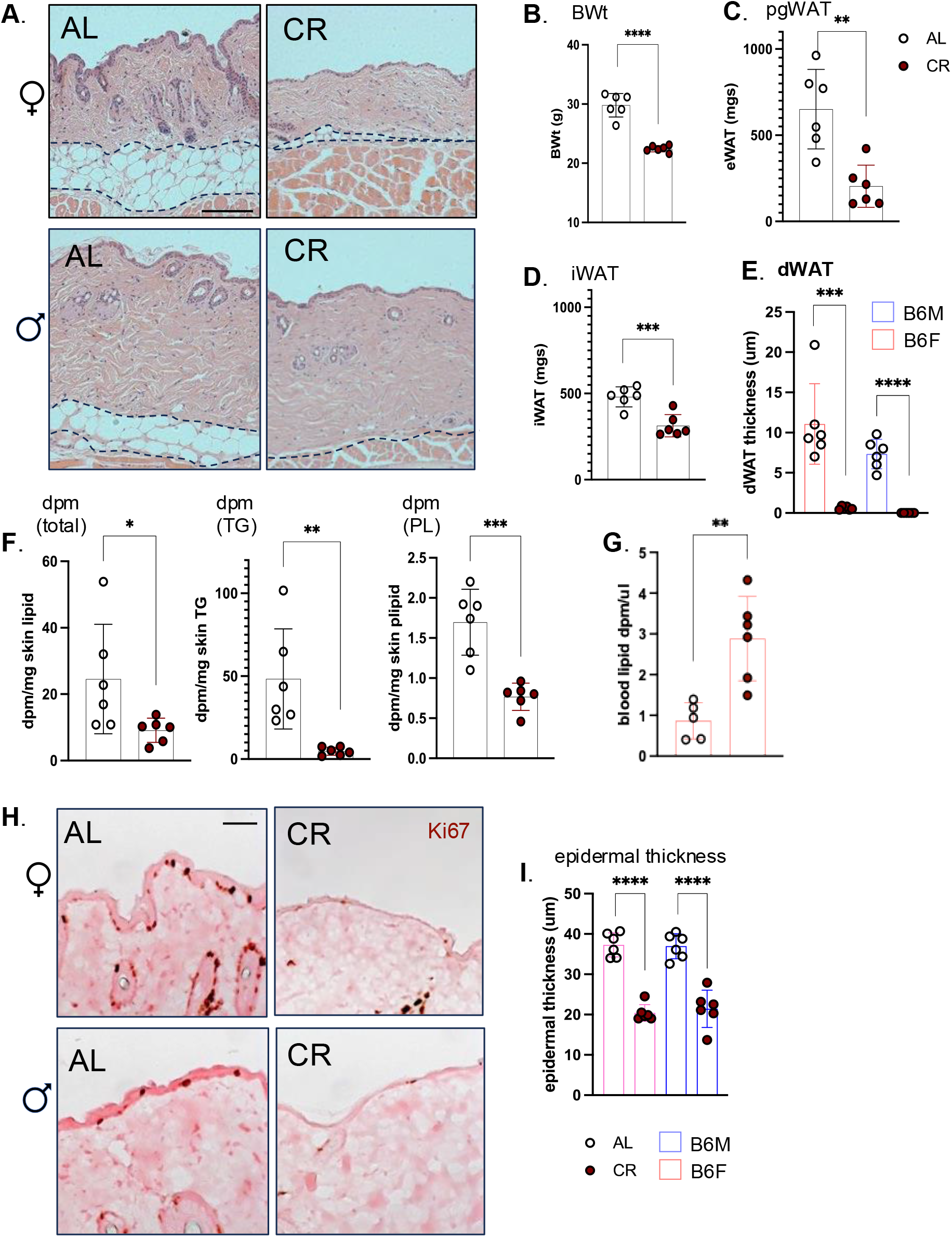
Skins of calorie restricted mice show minimal lipid uptake, depleted dWAT and a reduced mitotic index in the interfollicular epithelium. **A.** Representative H&E-stained sections from skins of C57BL/6J male and female mice fed *ad libitum* (**AL**) or calorie-restricted (30% for 3 weeks; **CR;** n=6), illustrate profound dWAT depletion (quantified in **E**). Scale bar=100μm. **B-E.** The relative effect of CR on body weight (B), and adipose depots (**C**, pgWAT; **D**, iWAT) for C57BL/6J males illustrates extreme lipodystrophy of dWAT depot (**E**). **F.** Uptake of ^3^H labeled radiotracer into total skin lipid, or triglyceride (TG) or phospholipid (PL) purified by TLC for C57BL/6J males (Fig.S9). **G**. Circulating levels of ^3^H labeled radiotracer. **H**. The mitotic index of keratinocytes was evaluated by Ki67 staining of paraffin-embedded skins, and the thickness of the epidermis quantified in **I, J,** for both C57BL/6J males and females. Scale bar =25μm.

### Tracing of dietary lipids using the MCFA content of milk fat

As an alternative approach to demonstrating that skin-associated lipids reflect dietary lipids directly, we fed mice with a Western diet (**WD**), enriched in the medium chain fatty acids (12:0,12:1,13:0, 13:1,14:0,14:1,15:0; **MCFA**s) typical of milk fat^17^. Analysis of diets using LC/QTOF-MS confirmed that MCFAs are absent from standard chow, in which soybean oil (60%) and grains are the fat sources (**Fig.S3**).

Sera from mice fed WD for only 3 days showed typical changes, including increases in ceramides (CER), triglycerides (TG), phosphatidylcholines (PC), cholesterol esters (CE) other phospholipids (**Fig.5A**). Using LC/Q- TOF mass spectrometry, we identified TG species containing MCFAs in sera amongst the most most-changed group of lipids. MCFA-containing TGs were delivered to epidermis after feeding WD (**Fig.5B, C**), and the total amount of MCFA acyl chains significantly increased for dWAT, epidermis and sebome (**Fig.S4**).

**Fig. 5.**
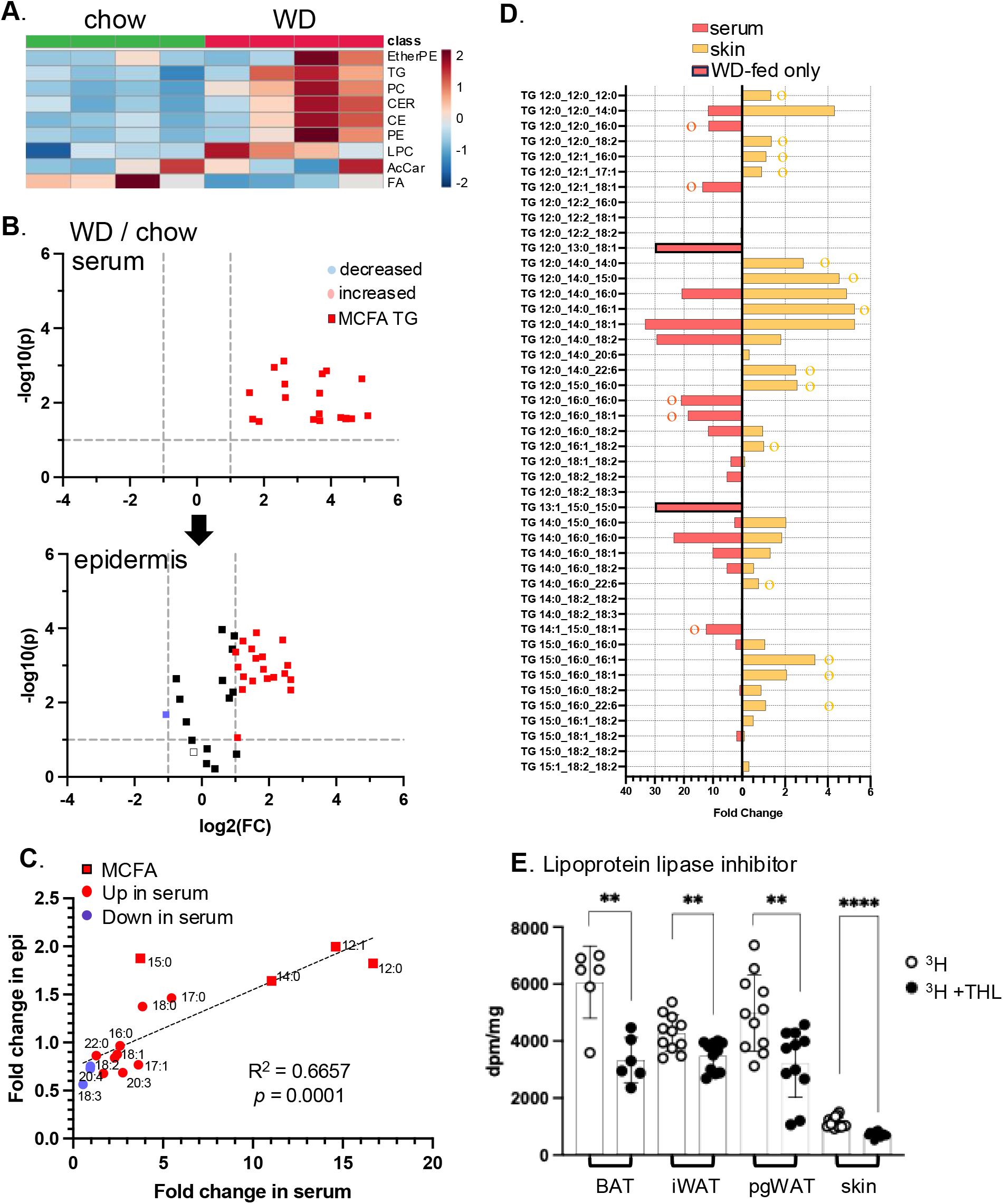
Dietary acyl chains are taken up by skin-associated triglycerides. **A.** Feeding of Western diet for 3 days induces typical changes of circulating lipid classes, shown here for C57BL/6J female mice (n=4; WD or chow-fed). Data is shown as an unsupervised heatmap of sera samples (EtherPE, ether phosphatidylethanolamines; PC, phosphatidylcholines; Cer, ceramides; CE cholesterol esters; PE, phosphatidylethanolamines; LPC, lysophosphatidylethanolamines; AcCar, acyl carnitines; FA, free fatty acids; n=4). Scale for heat map is shown as fold change. **B.** Volcano plots of the circulating lipidome and epidermis, showing significantly changed lipids (WD/chow), including TGs with medium chain fatty acids (MCFA), a signature of Western diet consumption. **C.** Fold change in specific acyl chains of serum and epidermis (epi) from 3d WD-fed mice is indicated, noting MCFA acyl chains. Total amounts of MCFA in serum, epidermis, sebome and dWAT of WD-fed mice is shown in Fig.S4. **D.** Fully specified MCFA-containing TGs were compared for serum and for skin, to test for a direct correlation of entire TG species (implicating direct delivery). Species marked with ο or o are unique to either serum or skin (respectively) or are present only in serum (WD-fed only). **E.** Specific activity of adipose depots and skins from C57BL/6J female mice administered tetrahydrolipstatin (THL, lipoprotein lipase inhibitor) before radiotracer administration (n=3). See also MCFA content and lipid profiles of diets (Fig.S3).

### An LPL-like enzyme is implicated in lipid uptake into skin

We considered whether dietary TG could be delivered intact to skin, seeking direct matches between increased circulating TGs with those observed in epidermis (skin; **Fig.5D**). We found many examples of MCFA-containing TGs that were increased in skin, but were not present, or did not increase, in the serum of WD-fed mice. This implicates a lipoprotein-lipase (**LPL**)-mediated acyl group reshuffling in the delivery of TG-derived acyl chains to the skin epithelial cell population. We administered the lipoprotein lipase inhibitor, tetrahydrolipstatin (**THL**), one hour prior to 3H-triolein feeding and showed that uptake into skin was significantly reduced, alongside delivery to other adipose depots, pgWAT, iWAT and BAT (**Fig.5E**). Therefore, LPL processing is likely to represent a rate determining step for the assimilation of TGs by skin.

### Within 3 days of feeding low-isoleucine WD, skins become more heat permeable

Given that our data show increased insulative properties of skins after high fat-diet feeding, we asked whether skin insulative properties could also respond to health-promoting diet interventions. We and others have shown that perhaps surprisingly, dietary protein restriction reduces fat mass and adiposity in both mice and humans^18–20^. Specifically, adiposity is promoted by branched-chain amino acids (BCAAs); restriction of all three BCAAs or isoleucine alone rapidly restores leanness to male mice with diet-induced obesity (DIO), associated with increased energy expenditure and better glycemic control^18,21^.

To demonstrate the effect of isoleucine restriction in female mice, we switched the diet of C57BL/6 female mice made obese by 16 weeks of WD consumption, to a diet with low isoleucine (**WDIL**) or low branched chain amino acids (leucine, isoleucine and valine, all at 33% of standard amounts, **WD 1/3xBCAA**; **Fig.6A,B**; **S5)**. The diets all contain identical amounts and sources of fat and sugar; calories removed through reductions of BCAAs are replaced with non-essential amino acids to maintain the diets as isocaloric (**Table S1**). We found that mice lost weight on WDIL diet, achieving a lean weight within 3 weeks (**Fig. 6A**). Analysis of body composition showed that the weight reduction in mice fed the BCAA and Ile-restricted diets primarily resulted from reduced fat mass (**Fig. 6B**, **S5**).

**Fig. 6.**
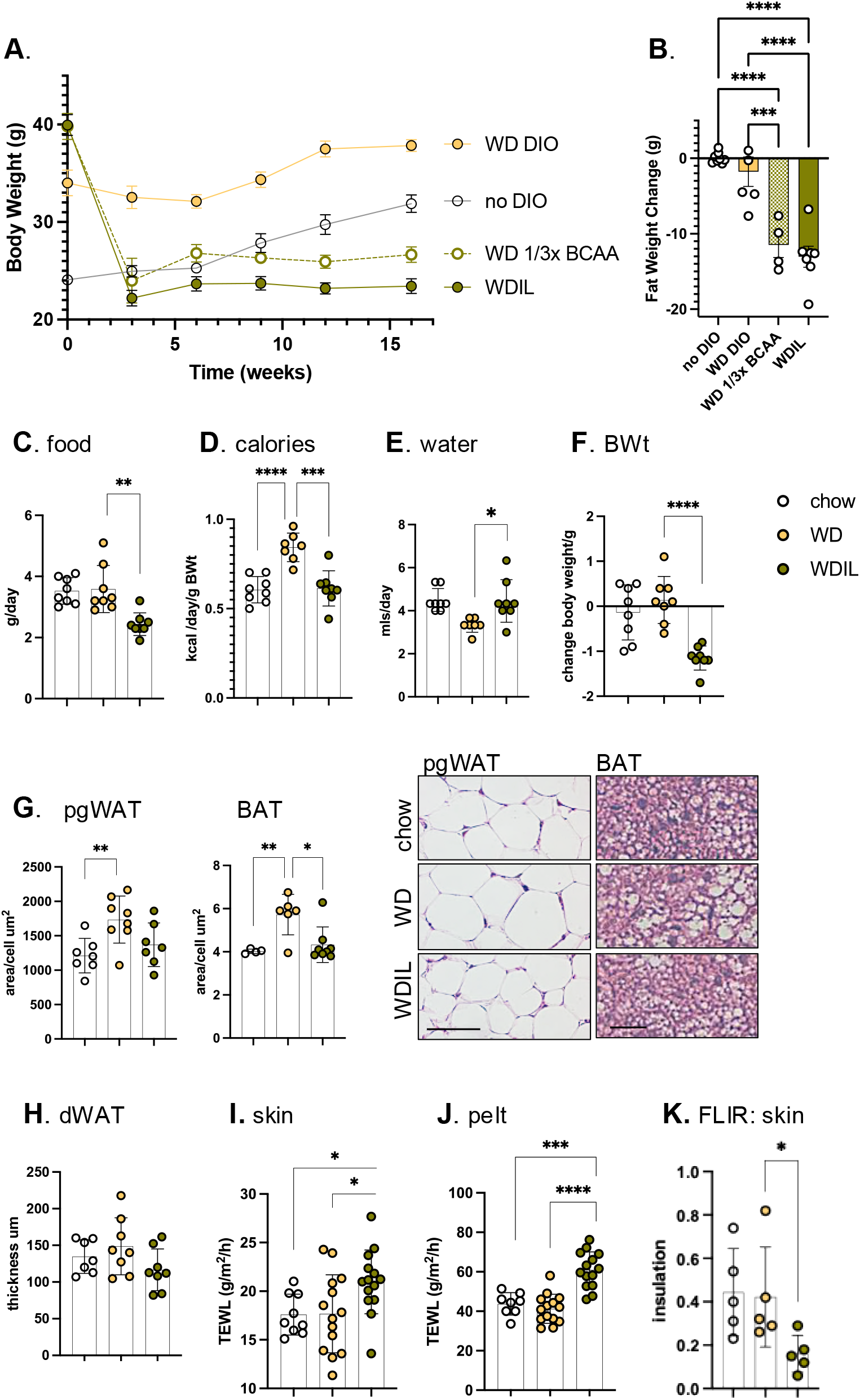
Mice fed a low-isoleucine Western diet rapidly develop heat-permeable skins. **A,B**. C57BL/6 female mice made obese by 16 weeks of WD feeding (**WD DIO;** TD.88137), were switched to low isoleucine (**WDIL**; TD.200692), low branched chain amino acid (**WD 1/3xBCAA**; TD.200691), or amino acid-defined WD (**WD DIO**; TD.200690). A control cohort were maintained on a regular fat content diet for the duration (**no DIO;** TD200693). B. Body composition was measured at 3 weeks post diet-switch (see also Fig. S5 for lean/fat body composition assay results; n≥4), illustrating the loss of nearly 15g of body fat within 21 days. **C**. To test the impact of a switch to WDIL diet feeding, female C57BL/6J mice (10-15 weeks old) were switched to Western diet (**WD**) or low- isoleucine Western diet (**WDIL)** for 3 days, and daily water and food consumption assessed (expressed as both grams/day or kcal/day/g body weight; **C-E,** n=7-8). **F**. Body weights were measured after 3 days**. G,H.** Changes to adipocyte depot lipid loads were calculated as the average area of pgWAT adipocytes, thickness of dWAT, or the lipid droplet area/cell for BAT, shown also as representative H&E-stained skin sections**. I-K.** TEWL and FLIR assays of thermal barrier function for mice fed chow, WD, or WDIL for 3 days (n=7-8, TEWL; n=5 FLIR).

**Fig. 7.**
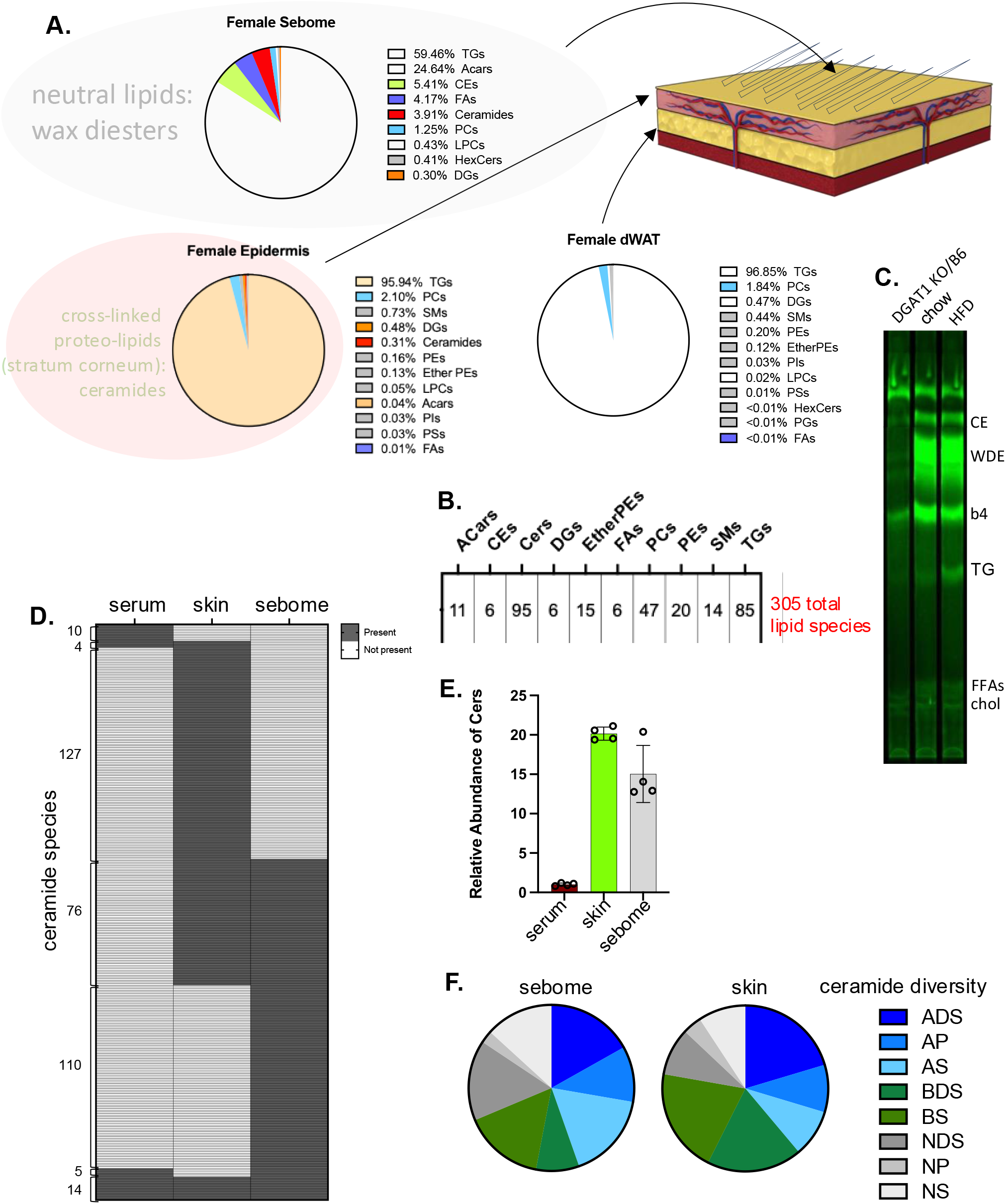
Multi-modal lipid analysis of skins. Skin tissue fractions produced as described in Fig.2 and Materials and Methods were analyzed for their lipid composition using LC/Q-TOF mass spectrometry. **A**. Using AUC as an approximation, the relative amounts of each detectable lipid class are described for each of sebome, epidermis and dWAT. The larger grey circle shown around the sebome indicates the neutral lipid wax esters (and potentially other classes) which are under-annotated by mass spectrometry. The larger pink circle around the epidermal lipids indicates the cross-linked proteolipid sheath called the stratum corneum, highly enriched with ceramides, which is not solubilized by these methods, therefore not detectable. **B.** The total number of reliably detected unique species identified for this analysis of epidermis or sebome from female mice fed HFD is shown. (Ceramide score includes both positive and negative modes). **C.** To interrogate the majority wax diester (WDE) class that predominates in sebome, WDE fractions are retrieved from preparative TLC plates and acyl chains derivatized to FAMEs, for analysis by GC/MS. Diacylglycerol O-acyltransferase (DGAT1) can serve as a wax synthase, and DGAT1 knockout mice have deficiencies in skin wax synthesis and function^74,75^, therefore extracts from mice with a DGAT1 mutation were used to indicate which fractions are likely to be wax diesters. CE, cholesterol esters; WDE, wax diesters: b4 (band4), unknown species; TG, triglycerides; FFAs, free fatty acids; chol, cholesterol. **D.** Comparison of ceramide species present in serum, skin and sebome (see also Fig.S6, annotated species), to illustrate the specificity of each ceramide cohort. **E.** Quantitation of total amounts of ceramide species identified by LC/QTOF-MS in serum, skin and sebome per mg of tissue (taken from positive mode data). **F.** Diversity of ceramide species shown for sebome and skin (total number of species identified in negative mode). Annotation key: S=sphingosine, P=phytosphingosine, DS=dihydro-sphingosine, A=α-hydroxy, B=β-hydroxy, N=non-hydroxylated.

To test whether loss of heat through skins could contribute to the rapid weight loss induced by this low-isoleucine diet, skin function was assessed for C57BL/6 female mice fed WD or WDIL for 3 days. Over this short timeline, mice fed WDIL adjusted their calorie intake to match chow-fed animals, where mice fed WD consumed 40% more calories (**Fig.6C,D**). Mice eating WDIL did not show the suppression of water intake induced by WD consumption (**Fig.6E**). After only 3 days of feeding, mice eating WDIL lost over 1 gram of body weight (**Fig.6F**). Mice eating Western diet showed no significant body weight increase at this time point, however, pgWAT adipocytes were already larger, and BAT adipocytes contained a higher lipid load (**Fig.6G**); mice eating a WDIL diet resisted these changes, and dWAT thickness was already trending down (**Fig.6H**).

Skins and pelts from WDIL-fed mice showed increased rates of evaporative cooling; TEWL was increased by 25% and 40% respectively (**Fig.6I,J**). The insulation measured by FLIR was reduced (**Fig.6K**). We conclude that within days of diet switch, WDIL-fed mice are protected from the obesogenic effects of WD-feeding and this is associated with the rapid development of heat-permeable skins. Note that in contrast to mice fed HFD, skin from mice fed WD show no net change in skin properties after 3 days feeding.

### Multimodal lipidomic analysis of dWAT, epidermis and sebome

We turned to molecular lipidomics to identify molecular correlates of altered thermal barriers. Each of the biomaterials comprising the skin thermal barrier depends upon different processes of lipid biosynthesis for their properties (**Fig.7A**). Specifically, TGs comprise most of the lipids of dermal WAT; the sebome is synthesized by sebocytes (ancillary to the hair follicle, and derived from epidermis) and comprises mostly wax diesters^22^ (shown by the TLC profile of **Fig.7C**). The epidermal stratum corneum is made by keratinocytes and comprises a cross-linked proteolipid matrix with abundant ceramides. We applied a LC/QTOF-MS untargeted lipidomics pipeline in both positive and negative modes, to identify and quantify unique lipid species^23^ (**Fig.7B**). We propose that neutral lipid species may be primarily responsible for thermal properties, however, the predominant wax diester (WDE) species are under-identified by standard lipid annotation programs. To report on this fraction, we separated WDE by TLC, derivatized the acyl groups to fatty acyl methyl esters (FAMEs), and characterized the products using GC/MS. One notable exception to the lipids assayed using this multi-modal approach is cholesterol, which is a major component of the stratum corneum^24,25^.

Serum, skin and sebome present almost exclusive ceramide profiles (**Fig.7D**,**S6**); this likely reflects the different functions of ceramides, as signaling molecules, precursors to the cross-linked ceramide sheath encasing mammals, and components of the thermal barrier coating mouse hair, respectively^2,26–29^. Skin has the most ceramide/mg tissue (**Fig.7E**), though hair/sebome shows similar abundance. Overall, most ceramide species in sebome and skin are hydroxylated (α- or β-) sphingosines and dihydro-sphingosines^30,31^ (**Fig.7F**).

### Three days of feeding HFD increases several lipid types in sebome and epidermis

Using separated skins from mice fed HFD for 3 days, we analyzed the lipid fraction of BALB/cJ male and female hair-associated sebome and epidermis (**Fig.8A, B**). Both epidermis and sebome were significantly changed by high fat consumption, characterized as a widespread depletion of approximately 20% of all lipids in the epidermis, and a corresponding gain of lipids secreted onto the pelt. Although the simplest explanation would be that the sebocytes empty their contents onto the pelt, we compared the lipids appearing in sebome with the lipids depleted in epidermis and found they were not the same. This suggests that epidermal lipid depletion is associated with a more generalized alteration of keratinocyte function^24^.

**Fig. 8.**
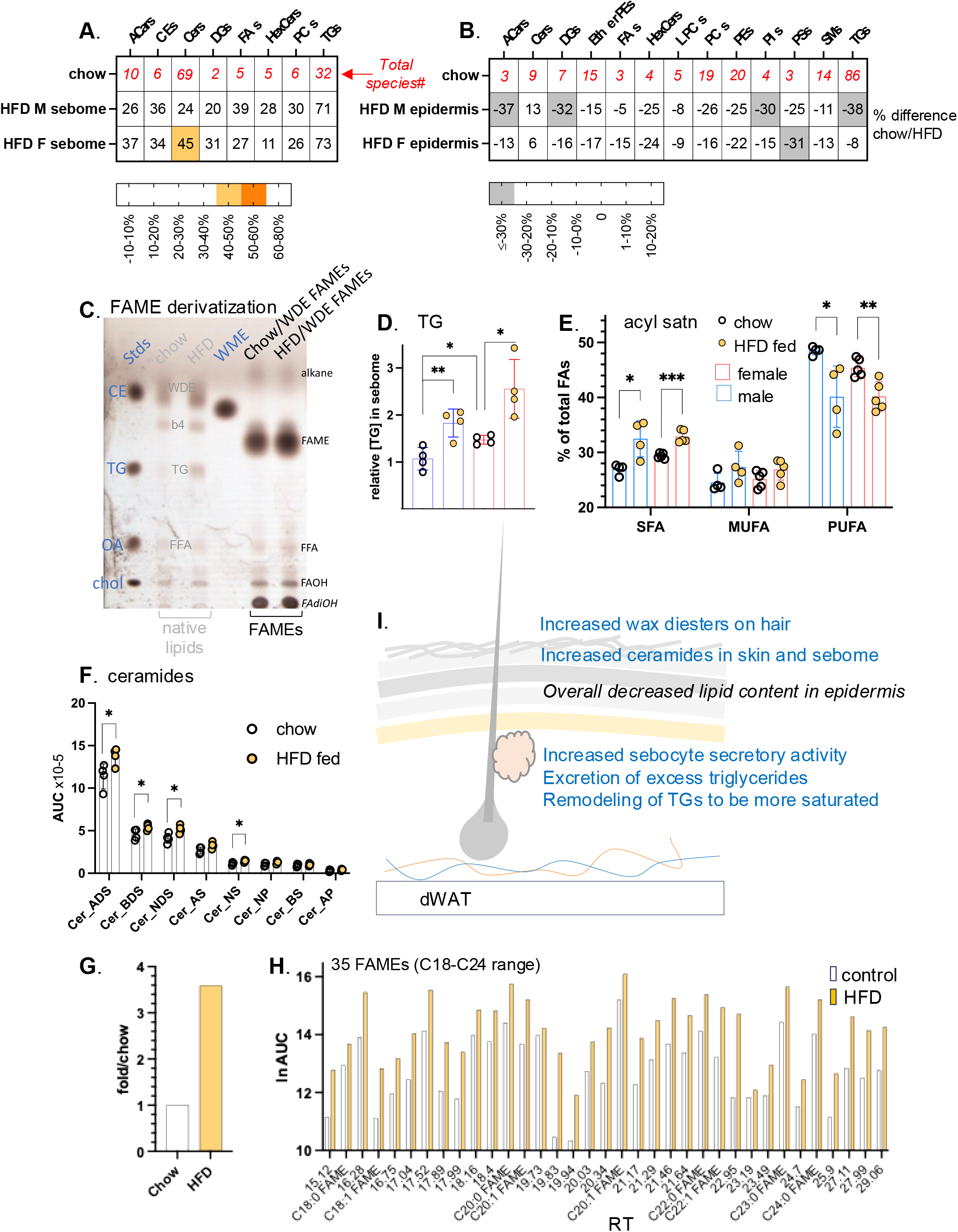
High fat diet consumption results in changes to components known to be thermally active in both sebome and skin lipidomes. **A,B.** The relative abundance (HFD/chow) of each lipid class is shown for sebome and epidermis from both BALB/cJ males and females. This HFD/chow ratio is calculated for males and females separately, since their skins are so different. The numbers in red along the top row show the total number of species compared for each class (n=4; additional lipid abbreviations noted here are DGs, diacylglycerols; HexCers, hexaceramides; PIs, phosphoinositides; PSs, phosphoserines; SMs, sphingomyelins). **C.** Evaluation of efficiency of FAME derivatization of WDE fraction. Shown is a charred analytical TLC separation of lipid extracts from hair of mice fed control or HFD diets for 3 days (pooled from 3 mice, repeated twice on separate samples). The lipid mix directly extracted from hair is shown labeled in grey, the WDE fraction after base- esterification to derivatize acyl-fatty acid methyl esters (FAMEs) are shown labeled in **black** (no residual bands at WDE moblity). Standards are shown in blue. WDE, wax diester; b4, unidentified band 4; FFA (free fatty acids) and TGs are identified by comigration with standards. Tentative identification of alkanes and fatty dihydroxy alcohols (FAdiOH) is also indicated. **D**. Quantitation of the increase in TGs present in sebome from male and female BALBc/J mice fed HFD, together with the relative acyl chain composition of the TGs (**E**). **F.** Each of the top 5 ceramide classes in sebome of HFD-fed male BALB/cJ mice increases by approximately the same degree (n=4). **G,H**. Quantitation of total FAMEs (**G**), together with specific data on 35 FAME species derived from the wax diester fraction of sebome, observed in the resolution range of the GC column. Peaks are reported as retention times, chain lengths C18-C26. Nine FAME species were tentatively identified by comparison with FAMEs in the standard mix (see also Fig.S7 for abundance waterfall plot). **I.** Summary scheme of the changes of sebome and epidermis observed in response to HFD feeding.

The TGs present in the hair fraction increased by 70% (both male and female) and were distinguished from the changes in other lipid classes by their larger accumulation (**Fig.8A, C-D**). This was observed by both lipidomics and TLC. Furthermore, saturated acyl chains became highly enriched in TG species, whilst poly-unsaturated acyl chains became depleted (**Fig.8E**). Thus, males fed HFD show an increase of 5.5% saturated fatty acids (from 27% total in control fed condition; SFAs), and a decrease of 8.4% (from 48.5% total; PUFAs), where females show a similar but muted trend. Given that increased acyl chain saturation changes the melt temperature and physical properties of TGs, this may contribute to significant functional change.

For the epidermis, the only lipid class that did not become depleted were the ceramides. Mass spectrometric assay can measure only the soluble ceramide pool, precursors to the assembly of the highly insoluble and cross- linked stratum corneum^32^. Thus we cannot quantify the amount of stratum corneum present, but can only speculate that given the precursor pool was increased, the amount of stratum corneum was likewise increased. Ceramides present in the sebome increased by approximately 20% across all 8 classes identified (**Fig.8F**). The changes of ceramide amount were higher for males than females, a pattern that correlated with the exacerbated pelt insulation for males fed HFD (Fig.1E). Baseline amounts of ceramides in females were >2-fold higher.

Surprisingly, even the wax diesters coating the pelt were affected by short exposure to HFD diet consumption. Hair goes through a month-long asynchronous process of growth and involution^33^, implying that it is stable and unchanging, however, these data suggest that the wax coating is labile. Thus, WDEs accumulate over 3-fold on the coats of mice fed HFD (**Fig. 8G**). FAME analysis of the acyl-groups present in sebome WDE lipids shows a diverse group of at least 35 species, some tentatively identified using co-migration with standard species (**Fig.8H**). This group is likely to include hydroxylated, very long and desaturated isomers of the more common standards. There is little change of composition induced by high fat feeding, instead there is an increase in WDE amount. There are 10 long chain acyl groups that comprise 90% of total (**Fig.S7**), however, the complexity of the assembled WDE class could be vast, given that each lipid includes at least 3 acyl groups. Thus we propose that the reduction of complexity by FAME derivatization is the method of choice, giving better insight than their partial identification and quantitation in LC/QTOF-MS spectra.

The changes we observed in skins of mice fed HFD for 3 days are summarized in the scheme of **Fig.8I**: increased wax diesters on hair, increased ceramides in skin, increased sebocyte activity, with excretion of excess triglycerides and remodeling of the secreted triglycerides to be more saturated.

### Three days of feeding low-isoleucine WD suppressed changes associated with WD, and suppressed key signaling lipids

By way of contrast, we analyzed the hyper heat-permeable skins from mice fed WDIL using the same techniques. We found that WDIL-fed mice partly resisted the increase in MCFA-containing circulating lipids that accompanies digestion of WD, and likewise, resisted delivery and accumulation of MCFAs in skins (**Fig.9A**, **Fig.S8**). Changes in circulating lipids were reduced or prevented in WDIL-fed mice, including the phosphatidylcholines (and associated lysophosphatidylcholines), ceramides, sphingomyelins, and cholesterol esters (**Fig.9B**0, leading us to conclude that low isoleucine-containing diet consumption leads to a profound alteration of the processing of WD by gut and/or liver.

**Fig. 9.**
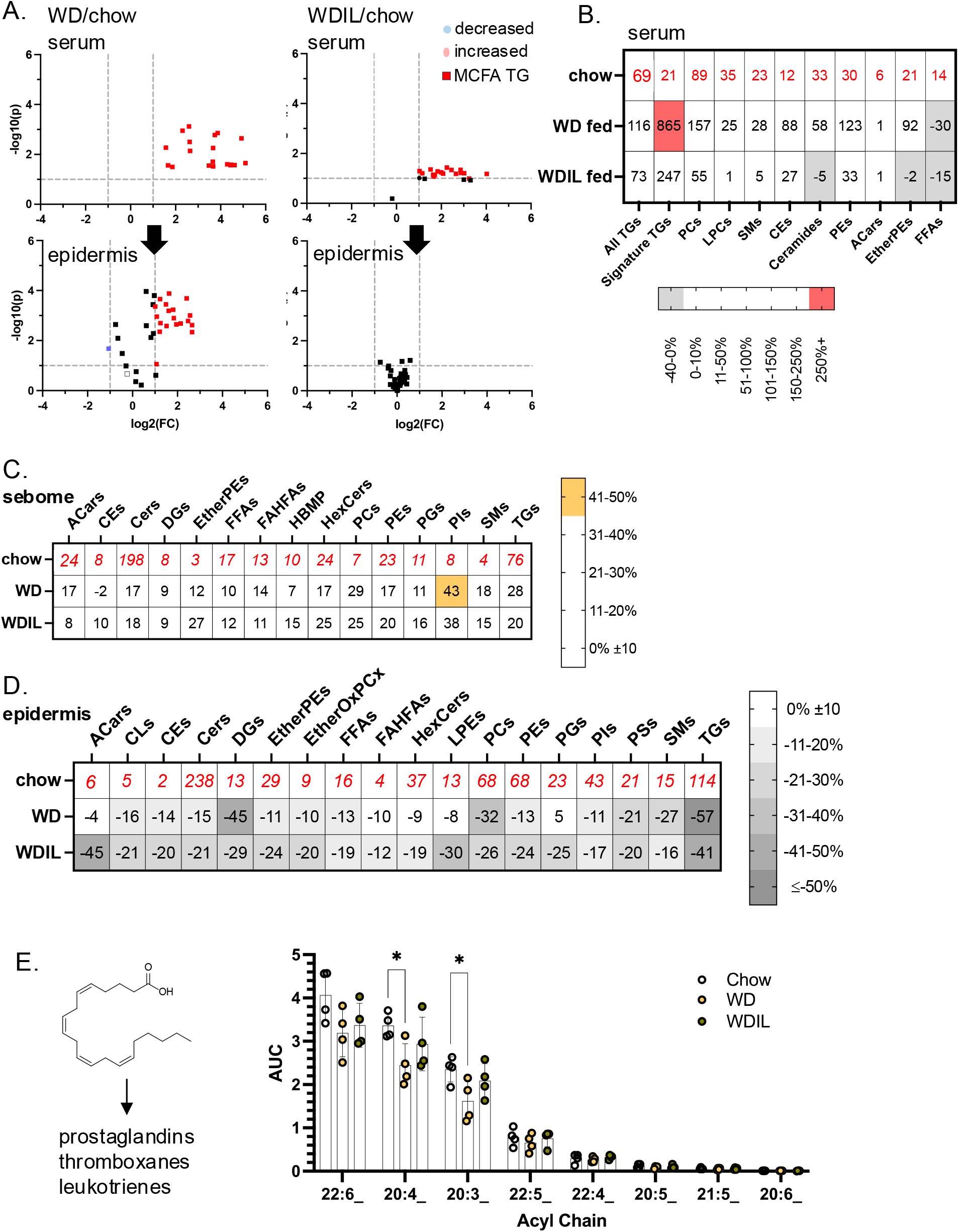
Low isoleucine Western diet prevents dietary delivery of acyl chains and mobilization of pro- inflammatory oxylipids from the epidermal reservoir. **A.** Relative appearance of Western-diet enriched MCFAs in epidermis of WDIL-fed C57BL/6J female mice compared to WD-fed mice (see also Fig.4; n=4). **B.** Heatmap of changes induced in circulating lipidome for WD- and WDIL-fed mice. **C,D.** The relative abundance of each lipid class is shown for C57BL/6J male sebome and epidermis, for control, WD and WDIL-fed mice. See also Fig.S8, inventory of total species interrogated. **E.** Oxylipid acyl chains, precursors of inflammatory cytokines (shown left hand side) were quantified for epidermal fractions from mice in all 3 diet conditions.

Similar to HFD-feeding, skins from WD-fed mice showed a broadly similar trend towards sebome lipid elevation with corresponding depletion of total lipids from epidermis. Interestingly, given that WDIL feeding appears to prevent most other changes, WDIL fed mice also show increased sebome. Unlike HFD consumption, the relative secretion of TGs was not increased above all the other lipid classes, therefore excess fat does not tend to vent out through the skins of WD-fed mice (**Fig.9C**; **S8**). For WD-fed mice, the epidermis-associated reservoir of oxylipid precursor acyl chains stored in TGs was significantly reduced (25%), notably 20:4 and 20:3 (**Fig.9E**); this change was prevented in mice fed WDIL. There were also distinctive changes characteristic of the epidermis of WDIL-fed mice, such as the depletion of acyl carnitines, phosphatidylethanolamines and related species of ether-linked phosphatidylethanolamines (Ether PEs), and ether-linked oxidized phosphatidylcholines (EtherOxPCs) (**Fig.9D**; **Fig.10A-D**). Acyl carnitines are key transporters of long chain fatty acyl chains during fatty acid oxidation^34^; PEs, Ether PEs and EtherOxPCs are peroxisomal in origin and are known to be important to lipid raft function^35^. Surprisingly, and in contrast to the response to HFD-feeding, WD-feeding reduced the amount of hair-associated wax diesters by nearly 80%, and mice fed WDIL-diet resisted this phenotype (**Fig.10E,F; Fig.S7**).

**Fig. 10.**
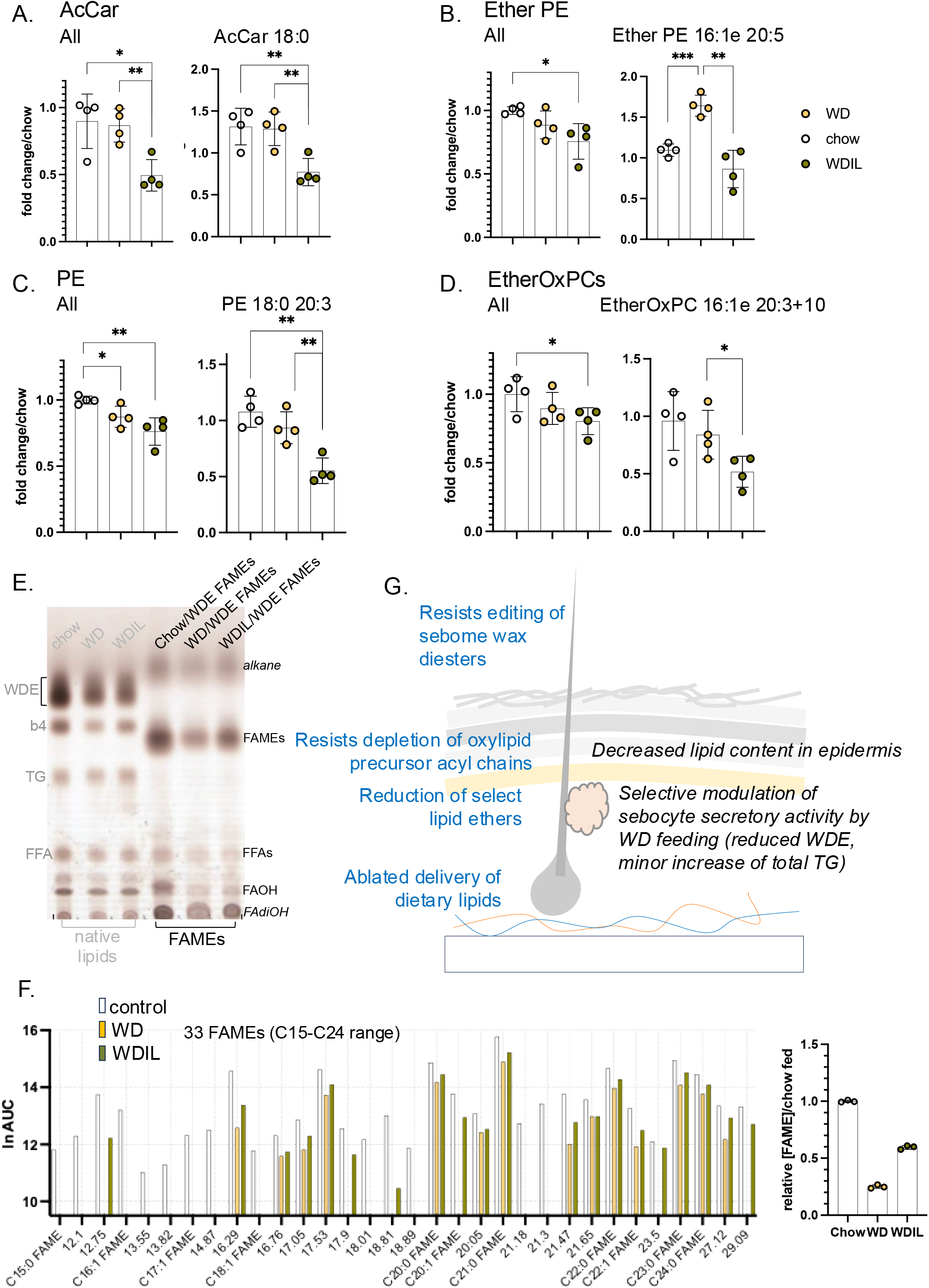
Summary of changes of heat-permeable skins induced by feeding low isoleucine diet. **A.** Several classes of lipids show changes specific to low isoleucine-fed C57BL/6J female mice (**A-D**, n=4). Illustrated are the trends present for the entire class, followed by a specific example of each. **E.** Quantitation of WDEs and FAME derivatization by TLC for all 3 diet conditions (annotated as for Fig. 8). **F.** FAME mixtures were analyzed by GC of WDE-derived FAMEs (prepared as for Fig.8H), illustrating the general suppression of WDE synthesis by WD consumption, and the reversal of this change when diet contains low isoleucine (see also Fig.S7, waterfall plot showing relative abundance). **G**. Summary scheme of the changes of sebome and epidermis observed in response to WD/WDIL feeding.

In summary, skins from WDIL-fed mice resist some, but not all, changes induced by WD-feeding, including delivery of dietary lipids, mobilization of oxylipid reserves and remodeling of wax diesters. However, this diet also induces novel responses, including reduced amounts of specific signaling lipids: together these molecular changes associate with increased skin heat permeability.

## DISCUSSION

### Skin is a major target for dietary lipids, and uptake is regulated

We have shown that skin is the dominant destination for triglyceride acyl chains, indeed more so than other individual tissues, including adipose and liver. Skin can store dietary fats for weeks after consumption; along with iWAT and BAT depots, skin shows a higher rate of mobilization and oxidation in thermogenically active mice (i.e. mice in sub-thermoneutral housing).

Triglyceride radiotracer was assimilated both by dWAT and by the epidermis, where the latter is known to rely upon TG stores for the massive biosynthesis reactions required to make the stratum corneum^24^. Furthermore, we show that calorie restricted mice show almost no uptake of ^3^H triolein by skin, despite high levels of circulating radiotracer, and we conclude that triglyceride uptake into skin is a regulated process. Lack of triglyceride uptake has profound consequences for the skins of calorie restricted mice, leading to a dramatic epidermal thinning, and depletion of adipocytes in dWAT. A published study from Kowaltowski and colleagues^36^ showed that after 6 months of calorie-restriction, mouse pelt became more dense and packed with guard hairs; this hair regrowth was supported by an increase in the activity of follicular stem cells. The surface temperature of the skin was lower, and indeed, the cold stress induced by shaving these mice forced them into muscle protein oxidation and metabolic distress. This implies that skin/pelt is a highly homeostatic organ, with means to compensate for excessive heat loss.

Given that skins become so highly lipid depleted by calorie restriction, it may not be surprising that calorie restricted rats show poor skin wound healing^37^. Subcutaneous and dermal WAT depots have been shown to be mobilized during wound repair in mice, and this mobilization is required to promote healing^38–40^. We speculate that poor lipid uptake into skins may underlie the deficiencies observed during wound healing of diabetic patients^41^, and propose that, like adipose, skin could have an important role in the uptake of other circulating metabolites, including other lipids, glucose and branched chain amino acids^42^.

Our data suggests that LPL is involved in uptake of dietary acyl groups into skin, along with other better-studied adipose depots^43^, but we anticipate there will be other processes of metabolite delivery to maintain the high level of *de novo* lipogenesis characteristic of sebocytes and keratinocytes.

It will be important to assess dietary lipid uptake into human skin. We have shown that there is little or no dWAT equivalent in human subjects, defined as a β-adrenergic-resistant depot associated with skin^44^. The regulation of skin-associated fat (SAF) in human may more closely resemble mouse scWAT. For lean women, we estimated SAF to be their largest adipose depot, varying between 6.5 −15 mm thick for individuals (an average of 15.8 kg) in a manner little related to general adiposity. Its regulation could be key to understanding personal energetics. Other studies of human adipose, specifically human visceral adipose tissue, showed slow TG turnover^45,46^, where turnover was regulated by β-adrenergic, insulin and other endocrine factors.

### How important is thermal adaptation of skin to insulation, energetics and metabolic health of mammals?

Our data shows that diet can induce rapid changes in the functional insulating properties of skin. Within only 3 days, high fat diet increases insulation (measured experimentally), and reduces heat lost by evaporative cooling by up to 28%, as measured for skins with or without hair, from both sexes of BALB/cJ and C57BL/6J mice. Since the consumption of high fat diet promotes skin-associated insulation, this could exacerbate the obesogenic effects of high fat consumption, by reducing the overall need for thermogenesis.

A prior publication made a sweeping conclusion that there was no change of insulation caused by obesity^47,48^ (with a counterpoint publication describing obese humans published by Brychta et al.^49,50^). Obese and normal weight mice were compared for their relative consumption of O2 at different environmental temperatures, and this was used to infer insulation based on the following assumptions: the rate of O2 consumption/joule of heat is constant, and the biological system is simple and non-adaptive. Since we know now that there are many BAT-independent heat production mechanisms (likely to have different stoichiometry of O2 consumption), and all manner of adaptive changes in obese mice, from body temperature to altered microbiome, it is timely to examine the specific factors which contribute to the total rate of energy expenditure^51^. By assaying skin function *ex vivo*, we eliminate the effect of β-adrenergic induced vasodilation, which is important when body temperature rises (this vasodilation is best illustrated by FLIR assays of the base of the tail^52^). Having said that, our study describes the molecular changes 3 days after diet switch. Since we have found that skin is uniquely flexible, homeostatic and adaptive (see example above), it will be important to characterize diet-induced changes long term, to test whether hyper-insulation is preserved.

### Which lipid components correlate with altered thermal barrier function?

Three days post diet-switch, there were already molecular changes in both the sebome and epidermis but none in dWAT: we suggest that dWAT acts as a more chronic adaptive response^53^. The observed gain of stratum corneum-associated ceramides^54^ and a trend to a more saturated acyl chain content are predicted to lower heat transfer. More novel perhaps are the changes we observed in the neutral wax diester classes which coat the skin and hair. Although hair only renews approximately monthly in an asynchronous pattern of replacement^5,55^, the wax diester coating was increased by >3-fold upon HFD-feeding. In contrast to this gain of insulation, feeding of a low isoleucine diet led to skins with increased heat permeability, associated with specific losses of lipids with signaling functions in the epidermal fraction. Taken together, these data suggest that thermal barrier function has contributions from several cell types.

### Which cellular components are implicated in functional responses?

Our study describes an acute modification of skin function: this is led by changes in the lipid products made by both keratinocytes and their differentiated derivatives, sebocytes. It is likely that both cell types are responsive and collaborative. Sebocytes have been called the brain of the skin^56^, due to their ability to integrate and process different cues, and they comprise some 2% of body mass. They are highly connected to neural and endocrine networks, have a high energy requirement, and their lipid products are important not only for hair shaft eruption but provide a signaling role to the skin cell community, providing feedback regulation of their own activity^57^.

### Does skin offer a means of excess lipid disposal?

Skin is highly regenerative and the ceramide component of stratum corneum is continuously shed. Ceramides are biosynthesized from triglyceride stores assembled by keratinocytes^24,28^, and together these could provide a significant route for elimination of excess lipid. Sebome secretion too provides a vent for excess dietary triglycerides, here increasing by 70% in mice fed high fat diet. We estimate that the whole pelt of a chow-fed mouse contains approximately 10 mg of TG, compared to 17 mg for a mouse fed HFD (assuming a total 200 mg hair per pelt). A pattern of triglyceride venting through sebome was suggested by Choa et al to be a major route for excess triglyceride secretion induced by the cytokine thymic stromal lymphopoietin (tslp)^58^. Although unlikely to account for the approximately 15g of adipose-associated triglycerides lost during the 3 weeks of this study, sebome-associated triglyceride levels could be an important biomarker for circulating triglyceride levels and/or changing energetics. Over-production of skin TGs was also noted after exposure to *acnes* bacteria^59^.

### Western and high fat diet promote different changes in skin

Skins from mice fed WD showed no net change in skin properties after 3 days feeding diet; this is associated with lack of accumulation of epidermal ceramide precursors and a dramatic reduction in sebome-associated WDE. These diets have different fat contents (calories from fat are 60% for HFD, 42% for WD and 18% for chow), however, WD combines high sucrose (34% w/w) with elevated fat content, a combination shown to be highly pro-inflammatory, both in general and for skin specifically^60–64^. For example, WD feeding exacerbated susceptibility to the TLR7 agonist imiquimod, leading to the development of psoriasiform dermatitis. A study of the molecular basis of balding induced by fish oil consumption revealed a clear direct relationship between dietary lipids, skin immune cells and their specific recruitment to the hair follicle^65^.

We found large reservoirs of oxylipid acyl precursor chains immobilized in the triglyceride fraction of epidermis, of which 25% are released during the first (3-day) exposure to WD consumption. These include 22:6, 20:4 and 20:3 acyl chains, all precursors to the inflammatory lipokines (prostaglandins, leukotrienes and thromboxanes). It is tempting to draw a conclusion that this mobilization provides the substrates for the triggered inflammatory reaction characteristic of skins from WD-fed mice.

### Skin changes in low-isoleucine fed mice reflect major changes in diet disposition

The functional changes of skins from mice fed WDIL diet coincided with a loss of about 1g of body weight over the course of 3 days, which we have previously shown to be the new, lean, steady state for these mice^18,21^. Sera from WDIL-fed mice revealed that low isoleucine can prevent the changes of circulating lipids that are typical of WD-consumption (elevation of phosphatidylcholines, sphingomyelins and ceramides). Likewise, uptake of dietary acyl chains to skin is prevented by low isoleucine feeding. However, low isoleucine diet does not merely prevent distribution of dietary lipids, the skins show a specific signature loss of lipids with signaling roles, including acyl carnitines, phosphatidylethanolamines and ether lipids such as ether phosphatidylethanolamines^66–68^, suggesting the induction of a specific signal.

Overall, we conclude that dietary lipids are taken up and stored by skins where they alter the properties of those skins, with potential to modulate animal insulation and energetics. We have shown that a “health promoting” diet is associated with increased heat loss through skins, whereas an obesogenic diet is insulating. Testing of other dietary paradigms will show how widespread this phenomenon is. Diet could modify the thermal properties of skins either directly, by incorporation into thermally-active lipids, or indirectly, via signaling to cells of the skin. The molecular changes documented here implicate diet as a modulator of both sebocyte and epidermal keratinocyte functions, where each is responsible for a different component of the thermal barrier.

## MATERIALS AND METHODS

### Ethical Approval. Mice

These studies were performed in strict accordance with the recommendations in the Guide for the Care and Use of Laboratory Animals of the National Institutes of Health. Experimental protocols were approved by the University of Wisconsin School of Medicine and Public Health Institutional Animal Care and Use Committee (IACUC) and the William S. Middleton Memorial Veterans Hospital IACUC. The number of mice used to perform this study was minimized, and every effort was made to reduce the chance of pain or suffering. All authors understand the ethical principles and confirm that this work complies with the animal ethics checklist.

### Mice

Mouse strains and sexes are specified in the results and figure legends: C57BL/6J (Jackson labs cat#00664) and BALB/cJ (Jackson labs cat#00651). For routine housing, mice were housed at constant temperature (19-23^0^C) in 12 h light/dark cycles with free access to water. The diets used were standard chow (Harlan Teklad Global Diet 2018), high fat diet (HFD; Envigo diet# TD.06414, 60% calories from fat); amino acid-defined Western diet (41% calories from fat, 21% w/w milk fat, 34% sucrose, Envigo diet# TD.160186) and a matching diet with low isoleucine (67% reduced (0.254%) isoleucine, Envigo diet# TD.170484), as described by Lamming and colleagues^18,21^. At the start of all diet switch experiments, mice were weight-matched and randomized to diet group. To demonstrate the long-term impact of low isoleucine diet consumption on mice with diet-induced obesity (DIO), C57BL/6J female mice were pre-conditioned by feeding a Western diet for 16 weeks (TD.88137), where the non-DIO cohort remained on a control diet (no DIO; TD.200693). Then WD-fed mice were divided between amino acid defined (AAD) Western diets containing low branched chain amino acids (WD 1/3x BCAA WD; TD.200691), only low isoleucine (WDIL; TD.200692), or standard amino acids (WD DIO; TD.200690). Full diet descriptions, compositions and item numbers are provided in Supplemental **Table 1**.

### Assay of skin thermal properties: transepidermal water loss (TEWL) and forward-looking infrared surface thermography (FLIR)

Half of the dorsal skin was shaved, and a strip of skin (approximately 3x2 cm) excised and assayed promptly for functional properties. Areas of skin obviously in anagen (appear black on underside for C57BL/6J skins) are not used for assay. Samples were transferred to a heat block protected from air flow, and held at 34^0^C; at least 7 TEWL readings were taken using a Vapometer probe (Delfin) of skin placed on a wet Wypall on the heat block, as previously described^6^. FLIR was measured using a FLIR One Pro camera plug-in for an iPhone, which is internally automatically calibrated. At least 10 pictures were taken to read out skin surface temperature together with near adjacent heat block temperature, and the difference read out as (arbitrary) ^0^C insulation.

### Calorie restriction, non-standard housing and metabolic phenotyping

For calorie restriction experiments, C57Bl/6J mice (both males and females; 16 weeks of age) were singly housed and fed ad libitum (**AL**) with control diet (D17110202; Research Diets) or calorie-restricted (30% **CR**, D19051601; Research Diets)^69^ for 3 weeks. The method used for evaluating radiotracer uptake had minor modifications **(Fig.S9).** For experiments with non-standard housing temperatures, mice were singly housed in environmental chambers (Memmert HPP750 or Caron 7350). Temperatures used were either thermoneutrality (**TMN;** 29^0^C) or cool (10^0^C), as described previously^51^. Many studies use thermoneutral temperatures >30^0^C; we use 29^0^C since we note no significant changes to breeding or exercise behavior at this housing temperature. We have not demonstrated that this temperature is thermoneutral for mice with acutely altered skin properties, however, the rate of thermogenesis (measured by BAT activation) is much reduced compared to the cool condition. For longer term experiments, mice were weighed weekly and body composition determined using an EchoMRI Body Composition Analyzer.

### Histological Analysis

Skin, BAT, perigonadal WAT (pgWAT) and the inguinal (mammary) subcutaneous fat pads (iWAT) were dissected for histological processing as described in Kasza et al ^44^. Briefly, paraformaldehyde-fixed, paraffin-embedded samples were H&E stained and assayed as follows: 1) dWAT thickness: 6 images of H&E-stained, non-anagen fields of skins (equivalent to ≥4500 μm linear dWAT) were assayed by image analysis (dividing total area by length). 2) Assay of BAT lipid stores: Lipid droplets were identified in gray scale images using circularity (0.1-1.0) and diameter (0.1-50 μm) thresholds in 6 independent fields of BAT (>1200 cells) and quantified using the open-source Fiji image processing package (https://loci.wisc.edu/software/fiji); data is expressed as average lipid droplet size (μm^2^). 3) pgWAT adipocyte size assay: 3-6 images across the length of the fat pad were scored for adipocyte size (measured as number of adipocytes/area scored) in each of 4-6 mice (total adipocytes scored >1500/mouse).

### Ki67 staining

The mitotic index of skins was determined using Ventana Discovery XT automated staining, as prescribed by the manufacturer, using the following antibodies: Abcam cat#ab16777 and HRP-conjugated anti-rabbit antibody (Ventana#760-4311). Slides were counterstained with eosin (0.3% in ethanol; Polysciences cat#02740).

### Radiotracer administration, assay and skin dissociation

Mice were acclimated to thermoneutrality before administration of Triolein [9,10-3H(N)] (ARC cat#0199), diluted 1:10 in canola oil. Mice were administered 10 μCi / 100 μl by gavage, and rehoused into controlled temperature housing, as indicated. Mice were euthanized and tissues dissolved in Solvable (Perkin Elmer cat#6NE9100) according to the manufacturer’s instructions, diluted into UltimaGold for liquid scintillation counting. Hair samples were collected for sebome analysis. For the preparation of separated skin fractions, skins were floated dermis side down onto cold elastase/DMEM/Hepes in a 6-well tissue culture dish for 6 hours. Tissue dissociation grade elastase (Sigma cat#E1250) was diluted to 0.12 mg/ml into DMEM (Gibco, high glucose cat#11965118) / 20 mM Hepes pH 7.4. Skins were rinsed in PBS, blotted, turned epidermis down and the dWAT layer scraped off manually with forceps. Purity of fractions was established by histology (**Fig.S10**). This method had minor modifications for the analysis of tissues from the calorie-restricted cohort (Suppl Method 1).

### Administration of lipoprotein lipase inhibitor, tetrahydrolipstatin (THL)

THL (Orlistat; Sigma cat# 04139)^70^ was dissolved in DMSO to 12.5 mg/ml, and diluted 10x prior to use in warm PBS. Drug was administered at 10 μg/g BWt by intraperitoneal injection, 1 hour prior to ^3^H triolein gavage, following the method of Bartelt et al^13^.

### Thin layer chromatography (TLC)

Lipids were extracted from hair (up to 50 mg) sequentially, using firstly 2:1 chloroform: methanol, and then acetone (4 mls each). Extracts were inverted to mix, incubated at room temperature, the extract decanted, combined and dried down. Lipids were resuspended in 4:1 chloroform:methanol, and loaded onto preparative TLC plates (Supelco TLC silica gel-60 glass plates; Millipore Sigma cat#1.00390.0001), and separated through 3 phases (as described by Choa et al^58^). The first phase was 80:20:1 hexane:isopropyl diether:acetic acid, to 50% plate height, second phase was 1:1 hexane:benzene to 80% plate height, and third phase was hexane, to 90% height. Lipids were visualized either using primuline (0.005% in 80:20 acetone:water)^13^ or charring with cupric sulfate (sprayed with 10% cupric sulfate in 8% phosphoric acid, dried, and baked at 300^0^F for 10 minutes).

### LC/QTOF-MS mass spectrometric analysis of lipids in tissues and sera

Lipidomic analysis was performed according to Simcox et al ^71,72^; briefly, 40μl aliquots of sera were combined with 250μls PBS and 225μls ice-cold MeOH containing internal standards (Avanti Splash cat#3307-07), and homogenized. Samples were then mixed with 750μls of ice-cold MTBE (methyl tert-butyl ether), rehomogenized, and separated by centrifugation (17,000g for 5 mins/4^0^C). The upper phase was transferred to a new tube, lyophilized and resuspended in 150 μls of isopropanol. Lipids were analyzed by UHPLC/MS/MS in positive and negative ion modes, at a dilution suitable to eliminate saturation issues. Extracts were separated on an Agilent 1260 Infinity II UHPLC system using an Acquity BEH C18 column (Waters 186009453; 1.7 μm 2.1 × 100 mm) maintained at 50 C with VanGuard BEH C18 precolumn (Waters 18003975), using the chromatography gradients described by Jain et al. The UHPLC system was connected to an Agilent 6546 Q-TOF MS dual AJS ESI mass spectrometer and run in both positive and negative modes as described. Samples were injected in a random order and scanned between 100 and 1,500 m/z. Tandem MS was performed at a fixed collision energy of 25 V. The injection volume was 2 μl for positive mode and 5 μl for negative mode.

### Lipidomic data processing

Methods for data processing were described by Jain et al^72^. Briefly, MS/MS data were analyzed using Agilent MassHunter Qualitative Analysis and LipidAnnotator for lipid identification^73^. Accuracy of retention times was checked by reference to internal standards. Data was imported into Agilent Profinder for lipid identification and peak integration (using sera-specific libraries). Data were analyzed using MetaboAnalyst free-ware (https://www.metaboanalyst.ca), Microsoft Excel and GraphPad Prism8 software (https://www.graphpad.com/scientific-software/prism).

A reporting overview is provided as Supplemental Data S1: *LIPIDOMICS DATASETS RESOURCES KEYS*. This includes a description of the total inventory screened (tab *total lipid number epi sebome*), and the inventory reported in Figures. The key to the data files provided in the massIVE repository, is provided, together with excel files supplemented with diagnostic ion species for each lipid, for each of 8 data sets in positive and negative acquisition mode, for serum, sebome, epidermis and dWAT.

### Fatty acid methyl ester (FAME) analysis

Lipids were scraped from TLC plates of separated lipids from 3 mice, and all analyses were performed in duplicate. Lipids were dissolved in toluene (100μl) and processed to fatty acid methyl esters by base-catalysed transesterification (Lipidmaps.org). Thus, 200μl of sodium methoxide (FisherSci cat#AC427228000) was added, incubated at 50°C for 10 mins, cooled and acidified with 10μl acetic acid. To the mixture, 0.5mls of water were added and vortexed, followed by 0.5ml hexane, vortexing and centrifugation to focus the interface. The top layer was removed, the hexane extraction repeated, and both supernatants pooled and dried down. Products of FAME reactions were evaluated by analytical TLC, run as for preparative TLC runs above, but using a hexane: diethyl ether: acetic acid (8:2:2) solvent run out for 10 cm, followed by charring with cupric sulfate. The FAME mixture was dissolved in anhydrous dicholoromethane, spiked with a C17 FAME standard (Cayman# 26723) and analyzed by GC/MS on an Agilent 7890B GC coupled to an Agilent 5977A MSD, with an Agilent DB-23 column (60m long, 0.25mm i.d., 0.25μm film thickness), with a staged injection protocol run at 250^0^C. Separating resolution, species identification and reproducibility were assessed using a Supelco 37-FAME standard (cat # CRM47885), and retention times (RTs) were compared to skin sample preparations.

### Statistical Analysis

The statistical tests appropriate to each analysis are indicated in Figure legends. To test for normal or lognormal distribution of sample values we used the Anderson-Darling test, outliers were identified using ROUT method. Box-and-whiskers graphs show median values with 10-90 percentile whiskers; other data are expressed as mean ± standard deviation, unless specifically stated.

## Acknowledgements.

We thank all the members of the Wisconsin Energy Expenditure Enthusiasts group together with Dr. Elizabeth Parks (MO), Dr. Dave Nelson (WI) and Dr. Bill Rizzo (NE) for their helpful discussion and providing experimental support. Additional assistance from Anu Singh from our TRIP lab was appreciated. The Lamming lab is supported in part by the NIA (AG056771, AG062328, AG081482, and AG084156) and the NIDDK (DK125859). M.E.T. is supported by F99AG083290. C-LEY is supported by RO1DK131752 and RO1DK124696. NR/IK/CMA were supported by a pilot from the Diabetes Research Center (DRC) at Washington University, P30 DK020579. RJ/JS are supported by R01DK133479, the UW-Madison Comprehensive Diabetes Center Core, the Office of the Vice Chancellor for Research and Graduate Education and Wisconsin Alumni Research Foundation, the University of Wisconsin-Madison School Department of Biochemistry, JDRF (JDRF201309442), and a Hatch Grant (WIS04000). JS is a HHMI Freeman Hrabowski Scholar and is an American Federation for Aging Research grant recipient. OAM is supported by RO1DK121759, RO1DK125513, RO1DK130879 and R01 AG069795.

## Declarations of conflicts of interest

DWL has received funding from, and is a scientific advisory board member of, Aeovian Pharmaceuticals, which seeks to develop novel, selective mTOR inhibitors for the treatment of various diseases.

## Author Contributions

Conceptualization, methodology: CMA, DL, RJ, JS. Experimental procedures: NR, CMA, IK, JM, GB-W, RJ. Resources: C-LEY, JS. Writing, editing: CMA, OM, DL.

## Funding sources

The Lamming lab is supported in part by the NIA (AG056771, AG062328, AG081482, and AG084156) and the NIDDK (DK125859). M.E.T. is supported by F99AG083290. C-LEY is supported by RO1DK131752 and RO1DK124696. NR/IK/CMA were supported by a pilot from the Diabetes Research Center (DRC) at Washington University, P30 DK020579. RJ/JS are supported by R01DK133479, the UW- Madison Comprehensive Diabetes Center Core, the Office of the Vice Chancellor for Research and Graduate Education and Wisconsin Alumni Research Foundation, the University of Wisconsin-Madison School Department of Biochemistry, JDRF (JDRF201309442), and a Hatch Grant (WIS04000). JS is a HHMI Freeman Hrabowski Scholar and is an American Federation for Aging Research grant recipient. OAM is supported by RO1DK121759, RO1DK125513, RO1DK130879 and R01 AG069795.

## SUPPLEMENTARY MATERIALS

**Supplemental Table**

Table S1. Diet composition

**Supplemental Data Files**

Data used to generate Figures (excel files Figs1-10)

## Supplemental Figures

#fig(accessory to Fig.1). Histology of skins from mice fed high fat diet (HFD) for 3 days Fig.S2 (accessory to Fig.3). Timeline of radiolabel accumulation with statistics.

#Fig.S3 (accessory to Figs.5,9). LC/QTOF-MS analysis of lipids in diets, to illustrate MCFA-TG amounts and other lipid species.

#Fig.S4 (accessory to Figs.5,9). Quantitation of MCFAs appearing in skin fractions, including dWAT. Fig.S5 (accessory to Figs.6,9,10). Changes of body composition in response to feeding of WDIL diet. Fig.S6 (accessory to Fig.7). Annotated listing of ceramides present in serum, epidermis and sebome.

#Fig.S7 (accessory to Fig.8, 10). Relative abundance of FAMEs in wax diesters from HFD and chow-fed sebome

#Fig.S8 (accessory to Fig.9). Total inventory of lipids identified during the analysis of C57BL/6J skins fed WD, WDIL or chow.

#Fig.S9 (accessory to Materials and Methods). Evaluation of radiotracer uptake into skins of calorie-restricted mice.

#Fig.S10 (accessory to Materials and Methods). Separation of dWAT from epidermis by elastase dissociation

**Data repository**. Lipidomic data is deposited at massIVE.ucsd.edu; submission# MSV000094882 (password CMA_052424!). Key to these data sets is included in the Supplemental Data files; *LIPIDOMICS DATASETS RESOURCES KEYS*.

## Notes

### Competing Interest Statement

The authors have declared no competing interest.

## REFERENCES

1 Radner, F. P., Grond, S., Haemmerle, G., Lass, A. & Zechner, R. Fat in the skin: Triacylglycerol metabolism in keratinocytes and its role in the development of neutral lipid storage disease. Dermatoendocrinol 3, 77–83 (2011). 10.4161/derm.3.2.15472

2 Kruse, V., Neess, D. & Faergeman, N. J. The Significance of Epidermal Lipid Metabolism in Whole-Body Physiology. Trends Endocrinol Metab 28, 669–683 (2017). 10.1016/j.tem.2017.06.001

3 Neess, D. et al. Epidermal Acyl-CoA-binding protein is indispensable for systemic energy homeostasis. Mol Metab 44, 101144 (2021). 10.1016/j.molmet.2020.101144

4 Neess, D. et al. Delayed hepatic adaptation to weaning in ACBP-/- mice is caused by disruption of the epidermal barrier. Cell Rep 5, 1403–1412 (2013). 10.1016/j.celrep.2013.11.010

5 Alexander, C. M. et al. Dermal white adipose tissue: a new component of the thermogenic response. J Lipid Res 56, 2061–2069 (2015). 10.1194/jlr.R062893

6 Kasza, I. et al. Evaporative cooling provides a major metabolic energy sink. Mol Metab 27, 47–61 (2019). 10.1016/j.molmet.2019.06.023

7 Sampath, H. & Ntambi, J. M. Role of stearoyl-CoA desaturase-1 in skin integrity and whole body energy balance. J Biol Chem 289, 2482–2488 (2014). 10.1074/jbc.R113.516716

8 Butera, A. et al. ZFP750 affects the cutaneous barrier through regulating lipid metabolism. Sci Adv 9, eadg5423 (2023). 10.1126/sciadv.adg5423

9 Liakath-Ali, K. et al. Alkaline ceramidase 1 is essential for mammalian skin homeostasis and regulating whole-body energy expenditure. J Pathol 239, 374–383 (2016). 10.1002/path.4737

10 Zhang, Z. et al. Dermal adipose tissue has high plasticity and undergoes reversible dedifferentiation in mice. J Clin Invest 129, 5327–5342 (2019). 10.1172/JCI130239

11 Marsh, D. Structural and thermodynamic determinants of chain-melting transition temperatures for phospholipid and glycolipids membranes. Biochim Biophys Acta 1798, 40–51 (2010). 10.1016/j.bbamem.2009.10.010

12 Parks, B. W. et al. Genetic control of obesity and gut microbiota composition in response to high-fat, high-sucrose diet in mice. Cell Metab 17, 141–152 (2013). 10.1016/j.cmet.2012.12.007

13 Bartelt, A. et al. Brown adipose tissue activity controls triglyceride clearance. Nature medicine 17, 200–205 (2011). 10.1038/nm.2297

14 Bagchi, D. P. & MacDougald, O. A. Identification and Dissection of Diverse Mouse Adipose Depots. J Vis Exp (2019). 10.3791/59499

15 Balasubramanian, P., Howell, P. R. & Anderson, R. M. Aging and Caloric Restriction Research: A Biological Perspective With Translational Potential. EBioMedicine 21, 37–44 (2017). 10.1016/j.ebiom.2017.06.015

16 Anderson, R. M. & Weindruch, R. The caloric restriction paradigm: implications for healthy human aging. Am J Hum Biol 24, 101–106 (2012). 10.1002/ajhb.22243

17 Mansson, H. L. Fatty acids in bovine milk fat. Food Nutr Res 52 (2008). 10.3402/fnr.v52i0.1821

18 Cummings, N. E. et al. Restoration of metabolic health by decreased consumption of branched-chain amino acids. J Physiol 596, 623–645 (2018). 10.1113/JP275075

19 Fontana, L. et al. Decreased Consumption of Branched-Chain Amino Acids Improves Metabolic Health. Cell Rep 16, 520–530 (2016). 10.1016/j.celrep.2016.05.092

20 Ferraz-Bannitz, R. et al. Dietary Protein Restriction Improves Metabolic Dysfunction in Patients with Metabolic Syndrome in a Randomized, Controlled Trial. Nutrients 14 (2022). 10.3390/nu14132670

21 Yu, D. et al. The adverse metabolic effects of branched-chain amino acids are mediated by isoleucine and valine. Cell Metab 33, 905–922 e906 (2021). 10.1016/j.cmet.2021.03.025

22 Nikkari, T. Comparative chemistry of sebum. J Invest Dermatol 62, 257–267 (1974). 10.1111/1523-1747.ep12676800

23 Tumanov, S. & Kamphorst, J. J. Recent advances in expanding the coverage of the lipidome. Curr Opin Biotechnol 43, 127–133 (2017). 10.1016/j.copbio.2016.11.008

24 Radner, F. P. & Fischer, J. The important role of epidermal triacylglycerol metabolism for maintenance of the skin permeability barrier function. Biochim Biophys Acta 1841, 409–415 (2014). 10.1016/j.bbalip.2013.07.013

25 Feingold, K. R. & Elias, P. M. Role of lipids in the formation and maintenance of the cutaneous permeability barrier. Biochim Biophys Acta 1841, 280–294 (2014). 10.1016/j.bbalip.2013.11.007

26 Summers, S. A., Chaurasia, B. & Holland, W. L. Metabolic Messengers: ceramides. Nat Metab 1, 1051–1058 (2019). 10.1038/s42255-019-0134-8

27 Albeituni, S. & Stiban, J. Roles of Ceramides and Other Sphingolipids in Immune Cell Function and Inflammation. Adv Exp Med Biol 1161, 169–191 (2019). 10.1007/978-3-030-21735-8_15

28 Rabionet, M., Gorgas, K. & Sandhoff, R. Ceramide synthesis in the epidermis. Biochim Biophys Acta 1841, 422–434 (2014). 10.1016/j.bbalip.2013.08.011

29 Chaurasia, B. & Summers, S. A. Ceramides in Metabolism: Key Lipotoxic Players. Annu Rev Physiol 83, 303–330 (2021). 10.1146/annurev-physiol-031620-093815

30 Kawana, M., Miyamoto, M., Ohno, Y. & Kihara, A. Comparative profiling and comprehensive quantification of stratum corneum ceramides in humans and mice by LC/MS/MS. J Lipid Res 61, 884–895 (2020). 10.1194/jlr.RA120000671

31 Uche, L. E., Gooris, G. S., Bouwstra, J. A. & Beddoes, C. M. Barrier Capability of Skin Lipid Models: Effect of Ceramides and Free Fatty Acid Composition. Langmuir 35, 15376–15388 (2019). 10.1021/acs.langmuir.9b03029

32 van Smeden, J. & Bouwstra, J. A. Stratum Corneum Lipids: Their Role for the Skin Barrier Function in Healthy Subjects and Atopic Dermatitis Patients. Curr Probl Dermatol 49, 8–26 (2016). 10.1159/000441540

33 Plikus, M. V. & Chuong, C. M. Complex hair cycle domain patterns and regenerative hair waves in living rodents. The Journal of investigative dermatology 128, 1071–1080 (2008). 10.1038/sj.jid.5701180

34 Li, S., Gao, D. & Jiang, Y. Function, Detection and Alteration of Acylcarnitine Metabolism in Hepatocellular Carcinoma. Metabolites 9 (2019). 10.3390/metabo9020036

35 Dean, J. M. & Lodhi, I. J. Structural and functional roles of ether lipids. Protein Cell 9, 196–206 (2018). 10.1007/s13238-017-0423-5

36 Forni, M. F. et al. Caloric Restriction Promotes Structural and Metabolic Changes in the Skin. Cell Rep 20, 2678–2692 (2017). 10.1016/j.celrep.2017.08.052

37 Hunt, N. D. et al. Effect of calorie restriction and refeeding on skin wound healing in the rat. Age (Dordr) 34, 1453–1458 (2012). 10.1007/s11357-011-9321-6

38 Cai, J. et al. The browning and mobilization of subcutaneous white adipose tissue supports efficient skin repair. Cell Metab 36, 1287–1301 e1287 (2024). 10.1016/j.cmet.2024.05.005

39 Schmidt, B. A. & Horsley, V. Intradermal adipocytes mediate fibroblast recruitment during skin wound healing. Development 140, 1517–1527 (2013). 10.1242/dev.087593

40 Shook, B. A. et al. Dermal Adipocyte Lipolysis and Myofibroblast Conversion Are Required for Efficient Skin Repair. Cell Stem Cell 26, 880–895 e886 (2020). 10.1016/j.stem.2020.03.013

41 Chen, V. Y., Siegfried, L. G., Tomic-Canic, M., Stone, R. C. & Pastar, I. Cutaneous changes in diabetic patients: Primed for aberrant healing? Wound Repair Regen 31, 700–712 (2023). 10.1111/wrr.13108

42 Felix, J. B., Cox, A. R. & Hartig, S. M. Acetyl-CoA and Metabolite Fluxes Regulate White Adipose Tissue Expansion. Trends Endocrinol Metab 32, 320–332 (2021). 10.1016/j.tem.2021.02.008

43 Wu, S. A., Kersten, S. & Qi, L. Lipoprotein Lipase and Its Regulators: An Unfolding Story. Trends Endocrinol Metab 32, 48–61 (2021). 10.1016/j.tem.2020.11.005

44 Kasza, I. et al. Contrasting recruitment of skin-associated adipose depots during cold challenge of mouse and human. J Physiol (2021). 10.1113/JP280922

45 Morigny, P., Boucher, J., Arner, P. & Langin, D. Lipid and glucose metabolism in white adipocytes: pathways, dysfunction and therapeutics. Nat Rev Endocrinol 17, 276–295 (2021). 10.1038/s41574-021-00471-8

46 Arner, P. & Ryden, M. Human white adipose tissue: A highly dynamic metabolic organ. J Intern Med 291, 611–621 (2022). 10.1111/joim.13435

47 Fischer, A. W., Cannon, B. & Nedergaard, J. No insulating effect of obesity, neither in mice nor in humans. Am J Physiol Endocrinol Metab 317, E952–E953 (2019). 10.1152/ajpendo.00333.2019

48 Fischer, A. W., Csikasz, R., von Essen, G., Cannon, B. & Nedergaard, J. No insulating effect of obesity. Am J Physiol Endocrinol Metab, ajpendo 00093 02016 (2016). 10.1152/ajpendo.00093.2016

49 Brychta, R. J. et al. The thermoneutral zone in women takes an "arctic" shift compared to men. Proc Natl Acad Sci U S A 121, e2311116121 (2024). 10.1073/pnas.2311116121

50 Brychta, R. J. et al. Quantification of the Capacity for Cold-Induced Thermogenesis in Young Men With and Without Obesity. J Clin Endocrinol Metab 104, 4865–4878 (2019). 10.1210/jc.2019-00728

51 Kasza, I. et al. "Humanizing" mouse environments: Humidity, diurnal cycles and thermoneutrality. Biochimie 210, 82–98 (2023). 10.1016/j.biochi.2022.10.015

52 Gordon, C. J. The mouse thermoregulatory system: Its impact on translating biomedical data to humans. Physiol Behav 179, 55–66 (2017). 10.1016/j.physbeh.2017.05.026

53 Kasza, I. et al. Syndecan-1 is required to maintain intradermal fat and prevent cold stress. PLoS Genet 10, e1004514 (2014). 10.1371/journal.pgen.1004514

54 Coderch, L., Lopez, O., de la Maza, A. & Parra, J. L. Ceramides and skin function. Am J Clin Dermatol 4, 107–129 (2003). 10.2165/00128071-200304020-00004

55 Plikus, M. V. et al. Cyclic dermal BMP signalling regulates stem cell activation during hair regeneration. Nature 451, 340–344 (2008). 10.1038/nature06457

56 Zouboulis, C. C., Yoshida, G. J., Wu, Y., Xia, L. & Schneider, M. R. Sebaceous gland: Milestones of 30- year modelling research dedicated to the "brain of the skin". Exp Dermatol 29, 1069–1079 (2020). 10.1111/exd.14184

57 Zouboulis, C. C. et al. Sebaceous immunobiology - skin homeostasis, pathophysiology, coordination of innate immunity and inflammatory response and disease associations. Front Immunol 13, 1029818 (2022). 10.3389/fimmu.2022.1029818

58 Choa, R. et al. Thymic stromal lymphopoietin induces adipose loss through sebum hypersecretion. Science 373 (2021). 10.1126/science.abd2893

59 Almoughrabie, S. et al. Commensal Cutibacterium acnes induce epidermal lipid synthesis important for skin barrier function. Sci Adv 9, eadg6262 (2023). 10.1126/sciadv.adg6262

60 Christ, A. et al. Western Diet Triggers NLRP3-Dependent Innate Immune Reprogramming. Cell 172, 162–175 e114 (2018). 10.1016/j.cell.2017.12.013

61 Herbert, D. et al. High-Fat Diet Exacerbates Early Psoriatic Skin Inflammation Independent of Obesity: Saturated Fatty Acids as Key Players. J Invest Dermatol 138, 1999–2009 (2018). 10.1016/j.jid.2018.03.1522

62 Higashi, Y. et al. High-fat diet exacerbates imiquimod-induced psoriasis-like dermatitis in mice. Exp Dermatol 27, 178–184 (2018). 10.1111/exd.13484

63 Shi, Z. et al. Short-Term Western Diet Intake Promotes IL-23‒Mediated Skin and Joint Inflammation Accompanied by Changes to the Gut Microbiota in Mice. J Invest Dermatol 141, 1780–1791 (2021). 10.1016/j.jid.2020.11.032

64 Yu, S. et al. A Western Diet, but Not a High-Fat and Low-Sugar Diet, Predisposes Mice to Enhanced Susceptibility to Imiquimod-Induced Psoriasiform Dermatitis. J Invest Dermatol 139, 1404–1407 (2019). 10.1016/j.jid.2018.12.002

65 Hao, J. et al. Consumption of fish oil high-fat diet induces murine hair loss via epidermal fatty acid binding protein in skin macrophages. Cell Rep 41, 111804 (2022). 10.1016/j.celrep.2022.111804

66 Kleiboeker, B. & Lodhi, I. J. Peroxisomal regulation of energy homeostasis: Effect on obesity and related metabolic disorders. Mol Metab 65, 101577 (2022). 10.1016/j.molmet.2022.101577

67 Nakamura, M. T., Yudell, B. E. & Loor, J. J. Regulation of energy metabolism by long-chain fatty acids. Prog Lipid Res 53, 124–144 (2014). 10.1016/j.plipres.2013.12.001

68 McCoin, C. S., Knotts, T. A. & Adams, S. H. Acylcarnitines--old actors auditioning for new roles in metabolic physiology. Nat Rev Endocrinol 11, 617–625 (2015). 10.1038/nrendo.2015.129

69 Li, Z. et al. Lipolysis of bone marrow adipocytes is required to fuel bone and the marrow niche during energy deficits. elife 11 (2022). 10.7554/eLife.78496

70 Augustus, A. S., Kako, Y., Yagyu, H. & Goldberg, I. J. Routes of FA delivery to cardiac muscle: modulation of lipoprotein lipolysis alters uptake of TG-derived FA. Am J Physiol Endocrinol Metab 284, E331–339 (2003). 10.1152/ajpendo.00298.2002

71 Simcox, J. et al. Global Analysis of Plasma Lipids Identifies Liver-Derived Acylcarnitines as a Fuel Source for Brown Fat Thermogenesis. Cell Metab 26, 509–522 e506 (2017). 10.1016/j.cmet.2017.08.006

72 Jain, R., Wade, G., Ong, I., Chaurasia, B. & Simcox, J. Determination of tissue contributions to the circulating lipid pool in cold exposure via systematic assessment of lipid profiles. J Lipid Res 63, 100197 (2022). 10.1016/j.jlr.2022.100197

73 Horing, M. et al. Accurate quantification of lipid species affected by isobaric overlap in Fourier-transform mass spectrometry. J Lipid Res 62, 100050 (2021). 10.1016/j.jlr.2021.100050

74 Yen, C. L., Stone, S. J., Koliwad, S., Harris, C. & Farese, R. V., Jr. Thematic review series: glycerolipids. DGAT enzymes and triacylglycerol biosynthesis. J Lipid Res 49, 2283–2301 (2008). 10.1194/jlr.R800018-JLR200

75 Yen, C. L., Monetti, M., Burri, B. J. & Farese, R. V., Jr. The triacylglycerol synthesis enzyme DGAT1 also catalyzes the synthesis of diacylglycerols, waxes, and retinyl esters. J Lipid Res 46, 1502–1511 (2005). 10.1194/jlr.M500036-JLR200

