## Supplemental Figures for "Dietary lipid is largely deposited in skin and rapidly affects insulating properties"

**Fig.S1. High fat diet consumption rapidly results in lower heat loss through skins** (supplementary to Fig.1C). Regardless of age, sex or strain, skins show no significant differences in gross histology 3 days after diet switch to HFD (including dWAT, dermal and epidermal thickness, follicle and sebocyte number). Mice were 7-9 months of age (Note C57BL/6J accumulate dWAT (and other adipose depots) with age[1]); n=4-6.

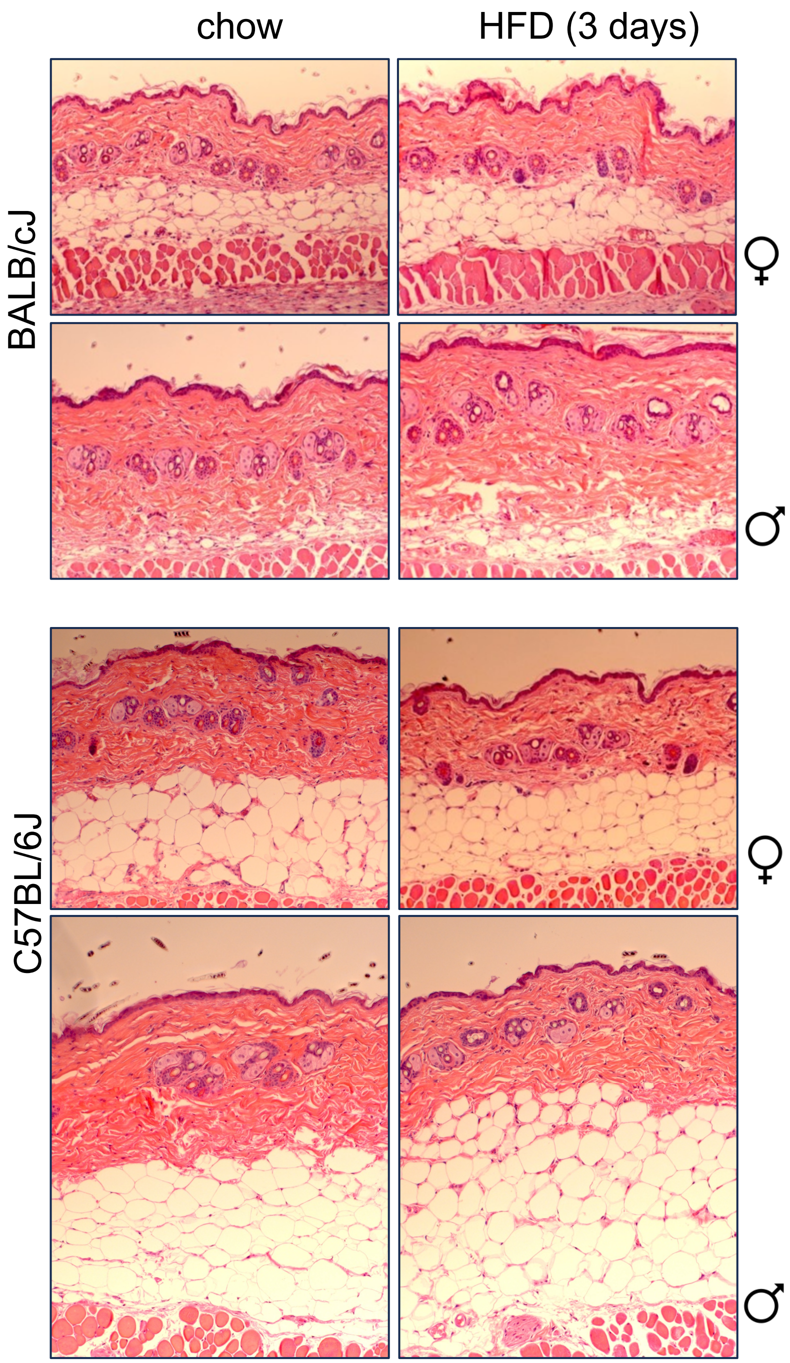


[1]Kasza, I., Hernando, D., Roldan-Alzate, A., Alexander, C. M. & Reeder, S. B. Thermogenic profiling using magnetic resonance imaging of dermal and other adipose tissues. *JCI Insight* **1**, e87146 (2016). https://doi.org:10.1172/jci.insight.87146

**Fig.S2** (accessory to Fig.3). **Dietary triglyceride radiolabel persists for 2 weeks in skin and adipose depots.** Average data shown in Fig.3 are shown here for independent mice, with statistical analysis.

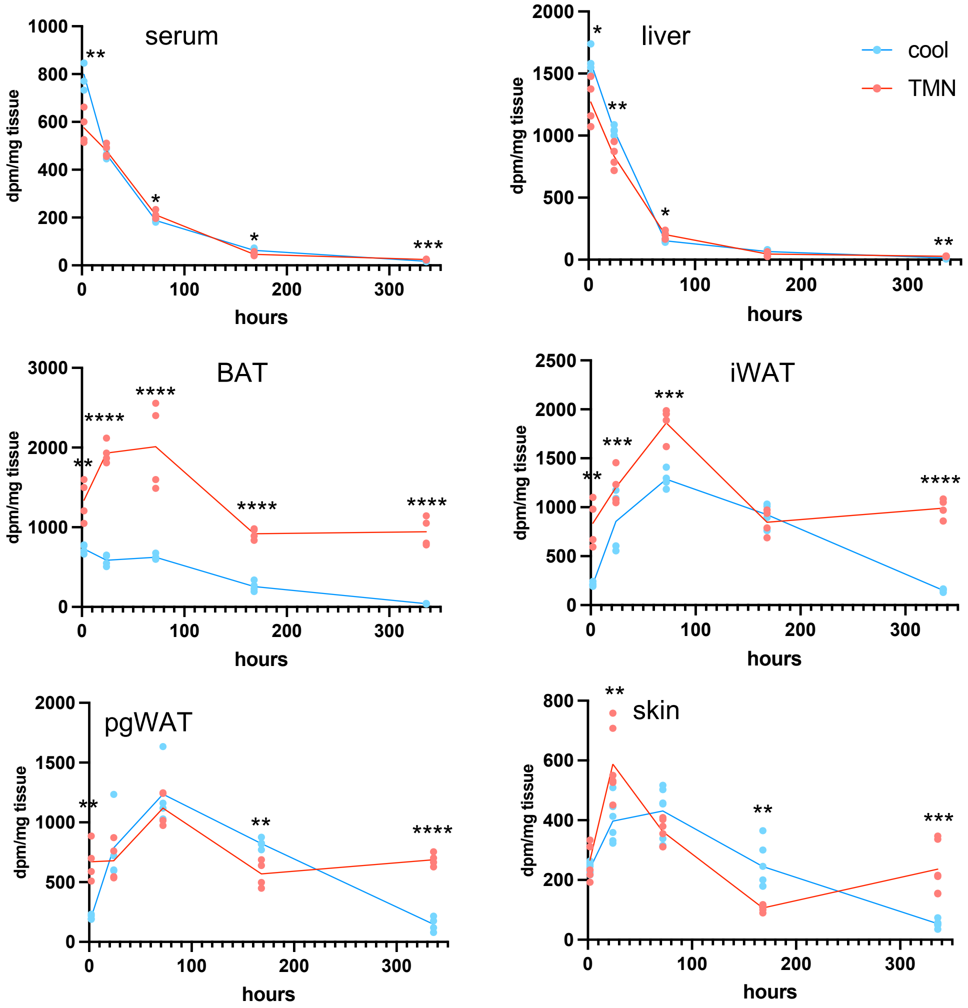

**Fig.S3** (accessory to Fig.5). **Milk fat acyl chains are delivered to skin-associated triglycerides.** WD, WDIL chow and HFD diets were analyzed using the same LC/QTOF-MS pipeline as the tissues, to characterize dietary lipids. Labels match Fig.5

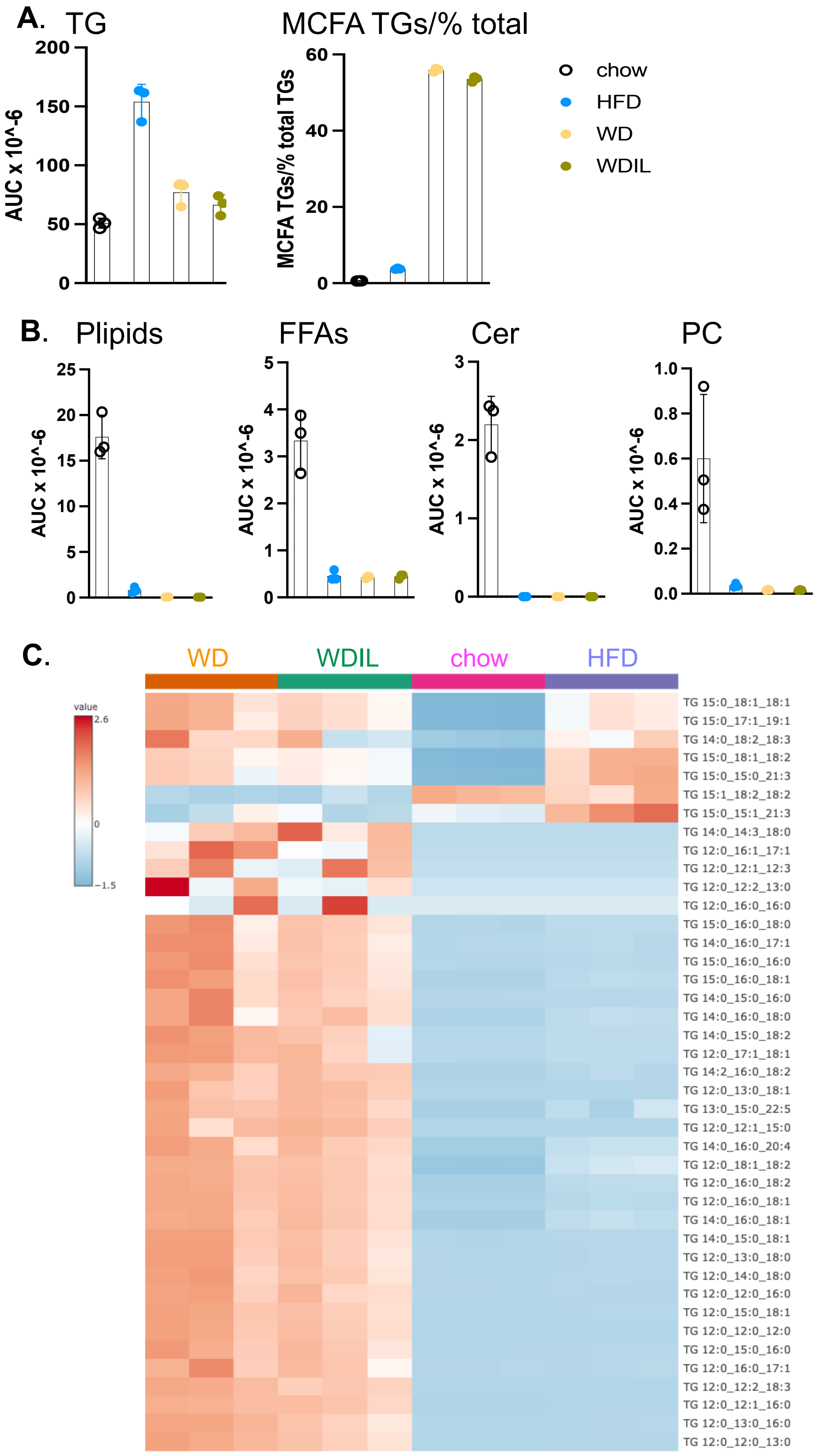

**Fig.S4** (supplementary to Fig 5, 9). **Milk fat acyl chains are delivered to skin-associated triglycerides**. **A**. Total amounts of MCFA acyl chain-containing triglycerides in dWAT collected from C57BL/6 female mice fed WD, WDIL for 3 days (compared to chow fed controls). **B**. Total amounts of specific MCFA acyl chains (in order of abundance), showing specific accumulation of 15:0 and 13:0 MCFAs. **C**. Fold change of MCFA acyl chains in serum, epidermis and sebome.

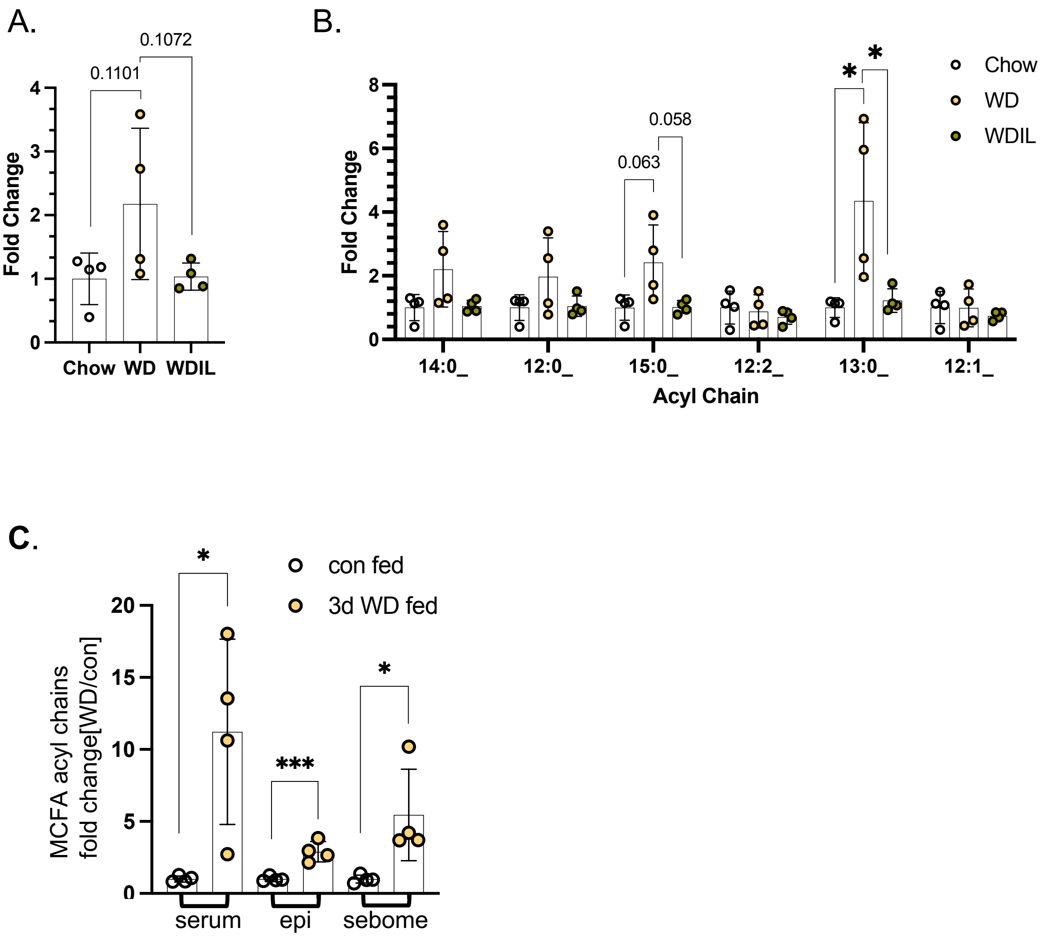

**Fig.S5. (**supplementary to Fig.6) **Mice fed a low-isoleucine Western diet rapidly develop heat-permeable skins.** C57BL/6 female mice were made obese by 16 weeks of WD feeding (WD DIO) or maintained at regular weight using a regular fat content diet (no DIO). Mice from the DIO cohort were then switched to low isoleucine (WDIL), low branched chain amino acid (WD 1/3xBCAA), or amino acid-defined WD (WD DIO). **A**. Lean and fat mass for each cohort, shown for a 16-week timecourse. A red arrow indicates the 3-week timepoint, statistically analyzed in **B:** Body composition of mice 3 weeks after diet switch, to illustrate statistical significance of changes after the diets have had their major effect.

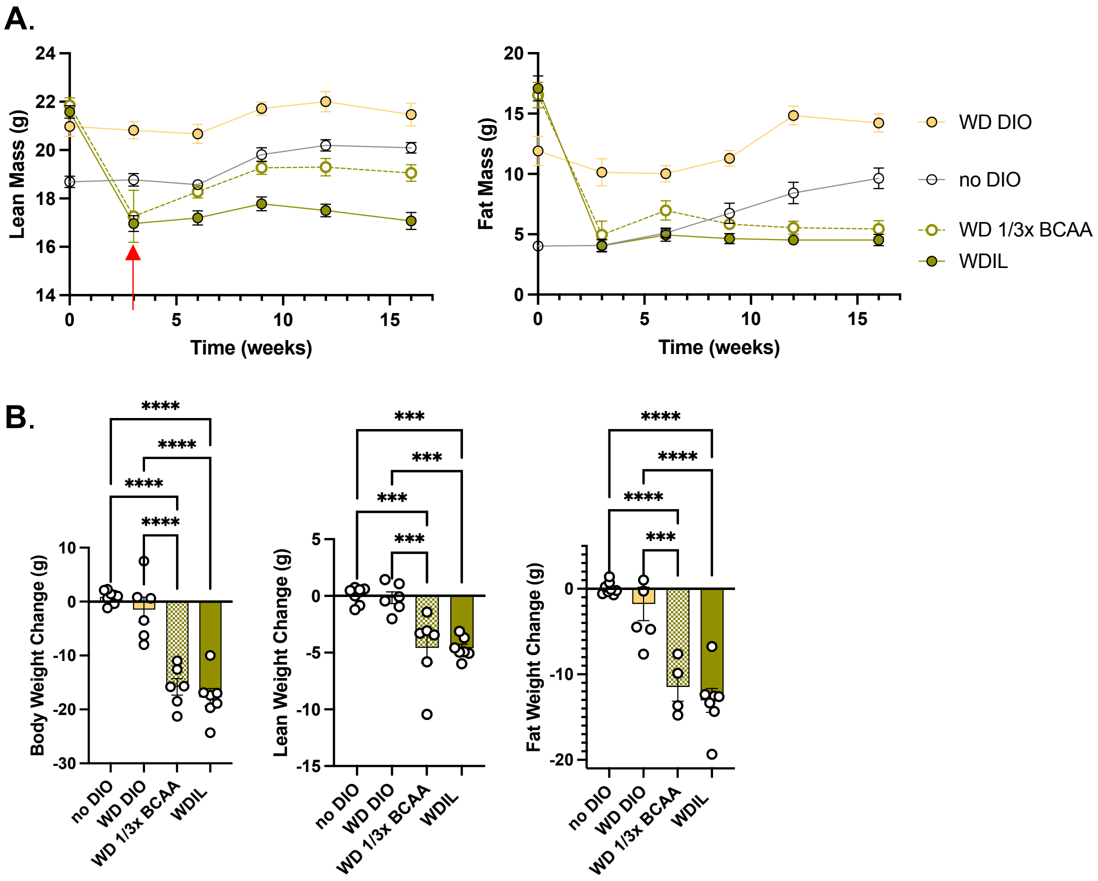


**Fig.S6. (next page;** supplementary to Fig.7) **Multi-modal lipid analysis of skins.**
Comparison of ceramide species present in serum, skin and sebome, with annotations

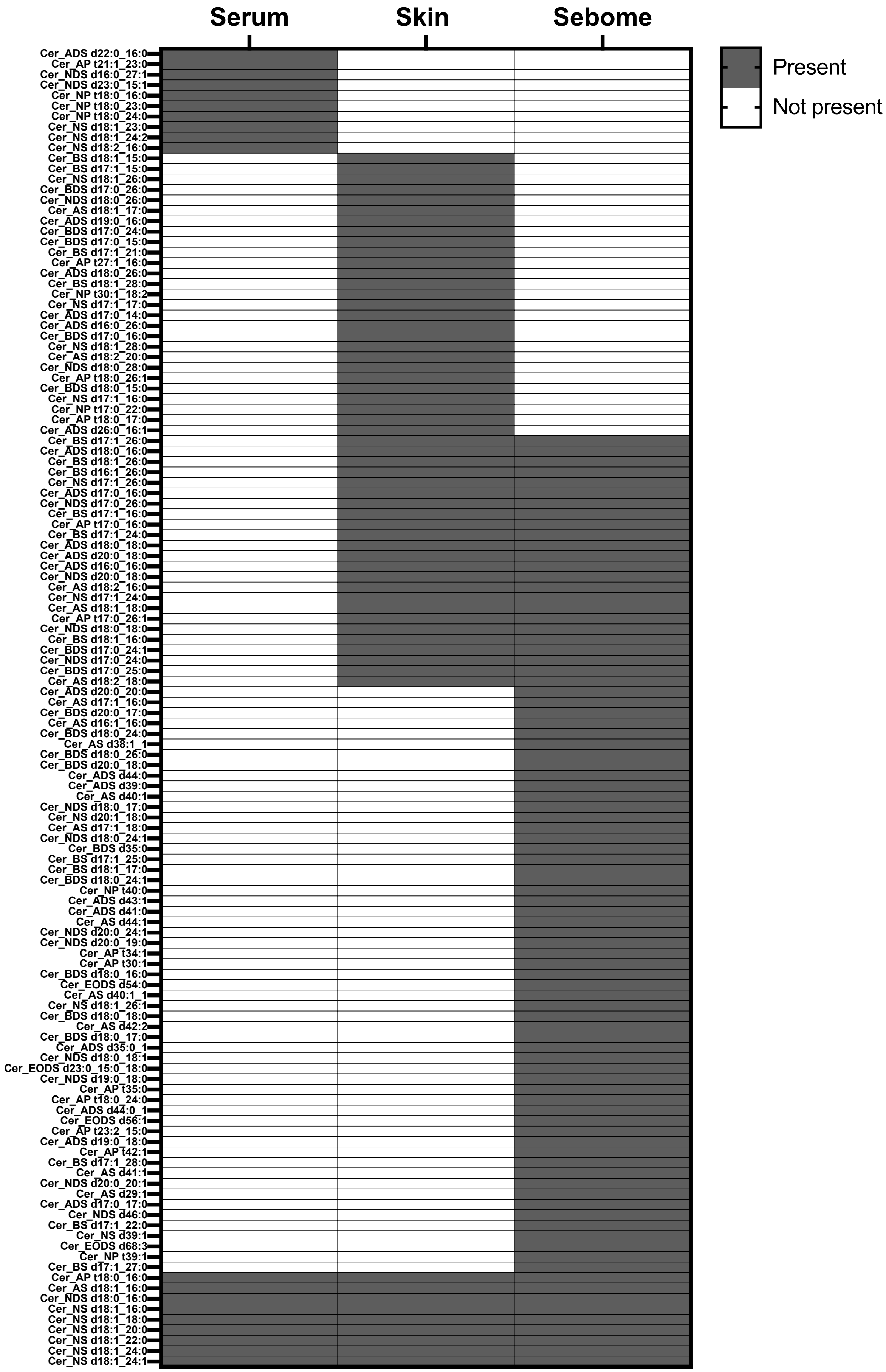

Serum
Skin
Sebome
Present
Not present
Cer_ADS d22:0_16:0
Cer_AP t21:1_23:0
Cer_NDS d16:0_27:1
Cer_NDS d23:0_15:1
Cer_NP t18:0_16:0
Cer_NP t18:0_23:0
Cer_NP t18:0_24:0
Cer_NS d18:1_23:0
Cer_NS d18:1_24:2
Cer_NS d18:2_16:0
Cer_BS d18:1_15:0
Cer_BS d17:1_15:0
Cer_NS d18:1_26:0
Cer_BDS d17:0_26:0
Cer_NDS d18:0_26:0
Cer_AS d18:1_17:0
Cer_ADS d19:0_16:0
Cer_BDS d17:0_24:0
Cer_BDS d17:0_15:0
Cer_BS d17:1_21:0
Cer_AP t27:1_16:0
Cer_ADS d18:0_26:0
Cer_BS d18:1_28:0
Cer_NP t30:1_18:2
Cer_NS d17:1_17:0
Cer_ADS d17:0_14:0
Cer_ADS d16:0_26:0
Cer_BDS d17:0_16:0
Cer_NS d18:1_28:0
Cer_AS d18:2_20:0
Cer_NDS d18:0_28:0
Cer_AP t18:0_26:1
Cer_BDS d18:0_15:0
Cer_NS d17:1_16:0
Cer_NP t17:0_22:0
Cer_AP t18:0_17:0
Cer_ADS d26:0_16:1
Cer_BS d17:1_26:0
Cer_ADS d18:0_16:0
Cer_BS d18:1_26:0
Cer_BS d16:1_26:0
Cer_NS d17:1_26:0
Cer_ADS d17:0_16:0
Cer_NDS d17:0_26:0
Cer_BS d17:1_16:0
Cer_AP t17:0_16:0
Cer_BS d17:1_24:0
Cer_ADS d18:0_18:0
Cer_ADS d20:0_18:0
Cer_ADS d16:0_16:0
Cer_NDS d20:0_18:0
Cer_AS d18:2_16:0
Cer_NS d17:1_24:0
Cer_AS d18:1_18:0
Cer_AP t17:0_26:1
Cer_NDS d18:0_18:0
Cer_BS d18:1_16:0
Cer_BDS d17:0_24:1
Cer_NDS d17:0_24:0
Cer_BDS d17:0_25:0
Cer_AS d18:2_18:0
Cer_ADS d20:0_20:0
Cer_AS d17:1_16:0
Cer_BDS d20:0_17:0
Cer_AS d16:1_16:0
Cer_BDS d18:0_24:0
Cer_AS d38:1_1
Cer_BDS d18:0_26:0
Cer_BDS d20:0_18:0
Cer_ADS d44:0
Cer_ADS d39:0
Cer_AS d40:1
Cer_NDS d18:0_17:0
Cer_NS d20:1_18:0
Cer_AS d17:1_18:0
Cer_NDS d18:0_24:1
Cer_BDS d35:0
Cer_BS d17:1_25:0
Cer_BS d18:1_17:0
Cer_BDS d18:0_24:1
Cer_NP t40:0
Cer_ADS d43:1
Cer_ADS d41:0
Cer_AS d44:1
Cer_NDS d20:0_24:1
Cer_NDS d20:0_19:0
Cer_AP t34:1
Cer_AP t30:1
Cer_BDS d18:0_16:0
Cer_EODS d54:0
Cer_AS d40:1_1
Cer_NS d18:1_26:1
Cer_BDS d18:0_18:0
Cer_AS d42:2
Cer_BDS d18:0_17:0
Cer_ADS d35:0_1
Cer_NDS d18:0_18:1
Cer_EODS d23:0_15:0_18:0
Cer_NDS d19:0_18:0
Cer_AP t35:0
Cer_AP t18:0_24:0
Cer_ADS d44:0_1
Cer_EODS d56:1
Cer_AP t23:2_15:0
Cer_ADS d19:0_18:0
Cer_AP t42:1
Cer_BS d17:1_28:0
Cer_AS d41:1
Cer_NDS d20:0_20:1
Cer_AS d29:1
Cer_ADS d17:0_17:0
Cer_NDS d46:0
Cer_BS d17:1_22:0
Cer_NS d39:1
Cer_EODS d68:3
Cer_NP t39:1
Cer_BS d17:1_27:0
Cer_AP t18:0_16:0
Cer_AS d18:1_16:0
Cer_NDS d18:0_16:0
Cer_NS d18:1_16:0
Cer_NS d18:1_18:0
Cer_NS d18:1_20:0
Cer_NS d18:1_22:0
Cer_NS d18:1_24:0
Cer_NS d18:1_24:1

**Fig.S7.** Supplemental to Fig.8, 10. **Analysis of acyl chains in WDEs of sebome for BALB/cJ HFD-fed and C57BL/6J WD/WDIL fed mice.** Abundance plots for FAMEs in WDE fractions. A. FAMEs from WDE of sebome from HFD and chow-fed mice. B. FAMEs from WDE of sebome from WD, WDIL and chow-fed mice

**A.**

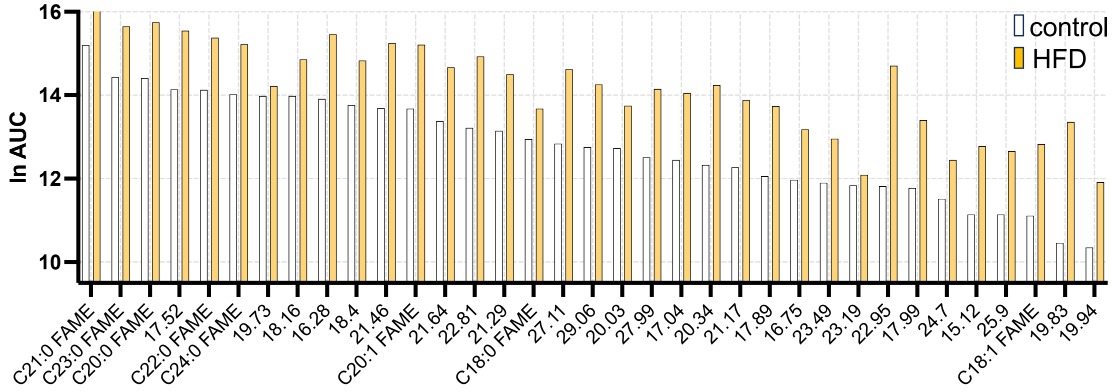


**B.**

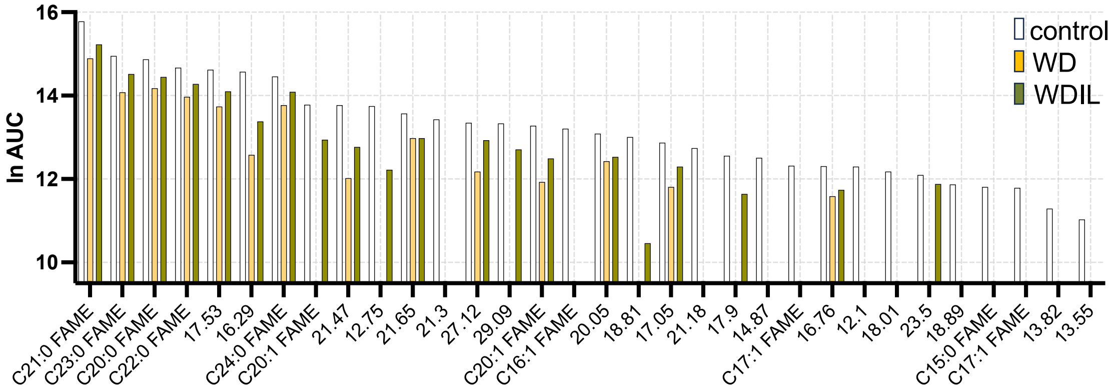

**Fig.S8.** Supplementary to Fig.9**. Low isoleucine Western diet prevents dietary delivery of acyl chains and mobilization of pro-inflammatory oxylipids from the epidermal reservoir.** The total inventory of lipids identified during the analysis of WD, WDIL and chow fed C57Bl/6J mice, to compare with Fig.7B

| ACars | CEs | Cers | DGs | EtherPEs | FAs | FAHFAs | PCs | PGs | PIs | PSs | SMs | TGs | |
|---|---|---|---|---|---|---|---|---|---|---|---|---|---|
| 23 | 9 | 273 | 17 | 31 | 21 | 14 | 39 | 27 | 33 | 12 | 24 | 142 | 665 total |

**Fig.S9.** Supplementary to Materials and Methods. **Evaluation of uptake of radiotracer into skins of calorie restricted mice.**

**Lipid extraction of tissues from calorie restricted cohort**

For reporting of 3H-triolein uptake into tissues of calorie restricted mice, lipids were extracted prior to counting. Tissues were pulverized, 25-50 mgs transferred to a glass homogenizer with 300 µls aq, and extracted with 1 ml chloroform:methanol (1:2), followed by transfer to a glass tube with 375 µls chloroform and 375 µls saline, and vortexing. Mixtures were separated by centrifugation at 2500 rpm for 10 mins, and the lower chloroform layer collected for analysis.

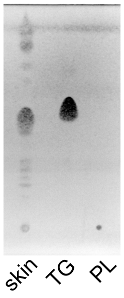


For reports of counts specifically associated with triglycerides and phospholipid fractions, extracts were separated by TLC and scraped off for evaluation.

**Fig.S10.** Supplementary to Materials and Methods. **Skin dissociation.** The purity of fractions obtained from elastase dissociation of dWAT from epidermis was established by histological analysis. These fractions (epidermis and dWAT) are enriched in the lipid-enriched fractions indicated, but include dermis and muscle respectively. Note the adipocytes collapse during extraction, and the fraction comprises extruded white fat

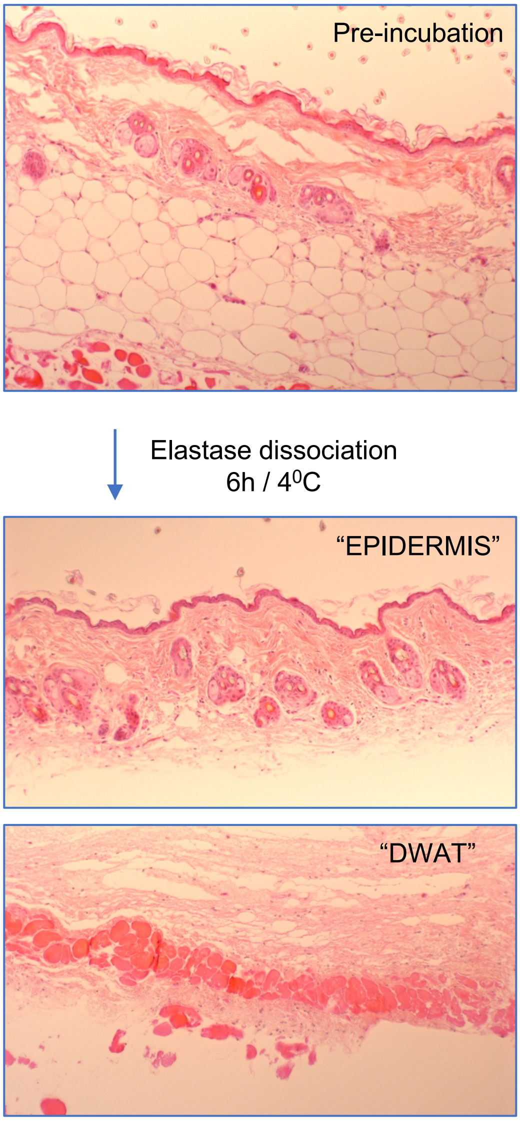
